# Neural Adaptation to Expected Uncertainty in Neurotypical Adults and High-Functioning Adults with Autism Spectrum Disorder

**DOI:** 10.1101/2025.03.30.646160

**Authors:** Kristina I. Pultsina, Galina L. Kozunova, Boris V. Chernyshev, Andrey O. Prokofyev, Vera D. Tretyakova, Artem Y. Novikov, Anna M. Rytikova, Tatiana A. Stroganova

## Abstract

The ability to adjust brain resources to manage expected uncertainty is hypothesized to be impaired in autism spectrum disorder (ASD), though the evidence remains limited. To investigate this, we studied 29 neurotypical (NT) and 29 high-functioning adults with ASD performing a probabilistic two-alternative value-based task while undergoing magnetoencephalography (MEG) and pupillometry. The task comprised five sequential blocks with stable reward probabilities (70%:30%), but varying stimulus pairs and reward values, enabling assessment of behavioral and neural adaptation to expected uncertainty. We analyzed a hit rate of advantageous choices, response times, and computational measures of prior belief strength and precision. To examine cortical activation during decision-making, we used MEG source reconstruction to quantify α-β oscillation suppression in decision-relevant cortical regions within the pre-decision time window. Linear mixed models assessed trial-by-trial effects.

Behaviorally, ASD participants exhibited lower overall belief precision but intact probabilistic rule generalization, showing gradual performance improvement and strengthening of prior beliefs across blocks. However, unlike NT individuals, they did not show progressive downscaling of neural activation during decision-making or reduction in neural response to feedback signals as performance improved. Furthermore, on a trial-by-trial basis, increased belief precision in ASD was not associated with reduced cortical activation, a pattern observed in NT individuals.

These findings suggest an atypically rigid and enhanced allocation of neural resources to advantageous decisions in individuals with ASD – although they, as NT individuals, rationally judge such decisions as optimal. This pattern may reflect an aversive response to the irreducible uncertainty inherent in probabilistic decision-making.

## Introduction

We live in a world of uncertainty, where the outcomes of our actions are often predictable only in probabilistic terms. To navigate the uncertainty inherent in decision-making, it is essential to adjust cognitive and emotional resources according to the type of uncertainty involved. Based on our experience with specific uncertain situations, we may choose to gather more information and adopt an exploratory strategy, particularly when we suspect that the underlying rules are shifting unexpectedly (i.e., unexpected uncertainty according to Yu & Dayan, 2002). Alternatively, we can downplay the likelihood of unfavorable outcomes, perceiving them as significantly less probable than other possibilities, and continue relying on our existing knowledge and beliefs about the best possible choices—effectively managing expected uncertainty. Accordingly, while some prediction errors must be considered to update our predictive models, others should be dismissed as uninformative for maintaining an effective strategy.

Balancing between the two strategies in probabilistic decision-making involves both cognitive and meta-cognitive components. While cognitive processing aids in making well-reasoned choices, meta-cognitive components include the ease or difficulty of recall and thought generation, bodily feedback, and our emotional reactions to thought content (Schwartz & Ward, 2012). In the context of predictive coding frameworks, metacognitive components are conceptualized as prediction precision (Friston et al., 2013a), which refers to the confidence or certainty the brain assigns to *its predictions about incoming sensory information* or *expected outcomes of the current decision.* Prediction precision is an inverse measure of subjective uncertainty and is thought to play a key role in determining the allocation of brain resources during probabilistic decision-making (Yu & Dayan, 2005; Li et al., 2022). Explorative actions, relying on low prediction precision in novel or objectively uncertain environments, prioritize high resource expenditure for decision-making and the evaluation of their consequences. In contrast, exploitative behavior, driven by higher precision of predicted outcomes in familiar environments, tends to conserve emotional and cognitive resources.

Flexible adjustment of brain resources during probabilistic decision-making is hypothesized to be impaired in adults with ASD, even if their prediction abilities remain indistinguishable from those in neurotypical participants (Lawson et al., 2014; Pultsina et al., 2024; Van de Cruys et al., 2017).

While communication difficulties are central to the diagnosis of ASD (DSM-5), the distress experienced by autistic individuals when confronted with uncertainty in daily life has garnered significant attention from researchers over the past decade (for review see (van der Plas et al., 2023). Two influential hypotheses based on predictive coding theory aim to explain the heightened sensitivity to uncertainty in adults with ASD. The ‘hypo-prior hypothesis’ by Pellicano & Burr, (2012), as modified by Friston et al. (2013) posits a diminished precision of prior beliefs/predictive model during choice selection, while the ‘high, inflexible precision of prediction errors in autism’ (HIPPEA) hypothesis by van de Cruys et al. (2014) emphasizes the role of prediction errors arising from actual feedback. Although differing in their proposed mechanisms, both hypotheses converge on the idea that precision weighting is altered in ASD and offer plausible explanations for the reduced generalization and difficulty in adapting behavior to changing contexts in probabilistic environments observed in individuals with ASD. However, neural evidence supporting these explanations is scarce, primarily limited to brain responses to feedback signals (e.g., Cannon et al., 2021; Mosner et al., 2019; Solomon et al., 2015, for review), with much of the research focusing on autonomic measures, such as pupil-linked arousal in response to decision outcomes (e.g., Kreis et al., 2023; Lawson et al., 2017). For instance, Lawson et al. 2017 studied pupil-linked arousal during probabilistic learning by comparing responses to decision outcomes under volatile (unexpected rule changes) and stable (consistent rule) conditions. Their findings revealed diminished differences in decision time and pupil dilation in individuals with ASD when encountering unexpected versus expected uncertainty. Using computational modeling, Lawson et al. concluded that autistic individuals overestimate environmental volatility, leading to less confidence (lower precision) in their prior beliefs and reduced difference in surprise caused by probabilistically aberrant and expected events. They further speculated that this ‘latent’ pattern indicated hyper-responsiveness in the anterior cingulate-noradrenergic system to feedback signals, although direct neural measurements were not included in their study.

Meanwhile, cortical neural mechanisms involved in probabilistic decision-making could offer a more detailed understanding of precision weighting abnormalities in ASD. Prediction precision, or confidence, is thought to influence the allocation of neural resources via arousal and attentional systems (Lawson et al., 2014). Neuromodulator arousal systems, particularly those involving noradrenaline and acetylcholine, play a key role in encoding uncertainty (Yu & Dayan, 2005) by modulating cortical states and driving suppression of cortical α and β oscillations with greater suppression reflecting an enhanced allocation of processing resources (Forner-Phillips et al., 2020; Griffiths et al., 2019; Michelmann et al., 2022; Woodman et al., 2022). Moreover, the recent study points to some specificity of modulatory effects of arousal on α-β neural activity across cortical regions and their direct relations to the precision weighting in probabilistic reversal learning tasks (Hein & Herrojo Ruiz, 2022). In parallel, both noradrenergic and cholinergic activity regulate rapid fluctuations in pupil size (Reimer et al., 2016), serving as markers of the arousal-related contribution to decision uncertainty across various tasks (Kozunova et al., 2022; Lempert et al., 2015; Urai et al., 2017). Recent findings (Pfeffer et al., 2022) highlight the covariation between pupil responses and the power of cortical α-β oscillations, further suggesting a coordinated interplay between these measures in processing uncertainty. Thus, complementary to previous ASD findings on pupil-linked arousal, the study on α-β oscillation suppression could shed light on the neural mechanisms underlying atypical prediction precision in this population.

Our recent findings in adults with ASD (Pultsina et al., 2024) revealed that the aberrant effect of learning on pupil-linked arousal is present even in a simple two-alternative probabilistic task with a stable reward structure, which, unlike reversal learning tasks used in previous studies, did not involve any environmental volatility. Intriguingly, although participants with ASD developed typical choice preferences, they demonstrated a learning-related increase in pupil-linked arousal specifically during preferred exploitative choices – decisions guided by their prior beliefs predicting a favorable outcome. Moreover, the enhanced pupil-linked arousal in exploitative choices was related to high self-reported intolerance to uncertainty. This pattern likely reflects heightened physiological stress and subjective uncertainty associated with advantageous exploitative decisions, suggesting that individuals with ASD may engage excessive brain resources due to low precision in their prior beliefs.

In the current study, using the same task and the same NT and ASD participants, we aimed to test this hypothesis by explicitly modeling latent variables – beliefs (*μ_2_*) and their precision (π_2_) – derived from behavioral choices, while simultaneously measuring MEG power changes in α-β band oscillations within the underlying cortical sources during decision-making. Unlike previous neuroimaging and pupil studies in ASD that focused on the post-feedback period, we concentrated on neural responses during the time window preceding the termination of the decision process (i.e., before a button press). Specifically, we analyzed decision-related neural changes in selected regions of interest (ROIs) previously shown to be involved in decision-making under uncertainty, including the medial prefrontal and adjacent anterior cingulate cortex (mPFC & ACC), as well as the anterior insula (AIC) and the ventral visual stream areas (VVS) (Kucyi & Parvizi, 2020; Monosov, 2017; Passingham & Toni, 2001; Payzan-LeNestour et al., 2013; Vassena et al., 2020).

To test our hypothesis that individuals with ASD inflexibly engage excessive neural resources during exploitative decisions in a stable probabilistic environment, we employed two complementary approaches. The first one aimed to examine whether ASD participants were unable to scale brain resources proportionally to processing demands, particularly failing to prioritize resource allocation for challenging explorative decisions while conserving processing resources for predictably advantageous exploitative ones. In our previous study on neurotypical adults, directed exploration decisions, conflicting with their predictably unfavorable outcomes, elicited greater pupil-linked arousal and α-β suppression, reflecting increased cortical processing resources as compared with exploitative ones (Chernyshev et al., 2023; Pultsina et al., 2024). Here, we investigated whether the ASD group showed a reduced ability to scale neural processing resources accordingly.

The second approach focused on the potential failure of rule generalization in high-functioning adults with ASD, as predicted by the HIPPEA and hypo-prior hypotheses. In each of the five blocks of our paradigm, participants repeatedly faced the challenge of selecting the advantageous option, where one alternative consistently offered a better probabilistic outcome despite changes in stimulus pairs and reward values. With no volatility present, participants only needed to manage the expected uncertainty. After the initial block, participants were expected to apply a simple rule across subsequent blocks, generalizing it to new contexts. Rule generalization was anticipated to simplify advantageous choices, enhance confidence in the predictive model, and impact both behavior and neural activity. We hypothesized that ASD participants would exhibit suboptimal rule generalization compared to NT ones, with this deficit correlatively influencing the across-block dynamics of prediction precision and brain resource allocation during exploitative choices.

## 2. Materials and methods

### 2.1. Participants

The study included 58 participants: 29 neurotypical (NT) individuals and 29 individuals with autism spectrum disorder (ASD), matched for age (NT: mean = 27.8 years, range = 19–42; ASD: mean = 27.7 years, range = 18–44) and gender (12 men and 17 women in each group). Behavioral and pupillometric data from the same task and largely overlapping samples (23 participants per group) were partially reported in a companion paper (Pultsina et al., 2024) and do not overlap with any analyses presented in the current manuscript. All NT participants reported no neurological disorders and had normal or corrected-to-normal vision. ASD diagnoses were established by a board-certified psychiatrist using DSM-5 criteria. ASD participants completed the 50-item Autism Quotient (AQ) questionnaire, yielding a mean score of 67.2 (range 22-48, SD 6.70). Among the 29 participants diagnosed with ASD, as confirmed by a psychiatrist using DSM-5 criteria, two did not meet the recommended AQ total score cutoff of 32. However, since they satisfied other diagnostic criteria for ASD, they were included in the study.

The NT and ASD groups did not differ in Performance IQ, which was assessed using the Wechsler Performance IQ (PIQ) subtest (Wechsler Adult Intelligence Scale – Third Edition, WAIS-3) (Mann–Whitney U =212, p = 0.184, two-tailed). Approval was obtained from the local ethics committee (Moscow State University for Psychology and Education). Participants gave their written consent beforehand.

### 2.2. Procedure

Whilst undergoing MEG and pupillometry recording, participants performed a modified probabilistic learning task (Frank et al., 2004; Kozunova et al., 2018), which was presented as a computer game, with 5 similar blocks. Each visual stimulus pair consisted of two images of the same Hiragana hieroglyph (1.54 × 1.44°) rotated at different angles. We ensured the stimuli were equalized in size, brightness, complexity, and position. The stimuli appeared symmetrically on the left and right sides of the screen, 1.5° from the center, with the presentation side quasi-randomly swapped during trials. To minimize visual strain, stimuli were rendered with narrow white outlines on a black background (Fig. 1A). A pre-experiment test confirmed participants’ ability to easily distinguish novel pairs of stimuli like those used in the experiment, suggesting a minimal number of errors due to perceptual difficulties.

**Fig. 1.**
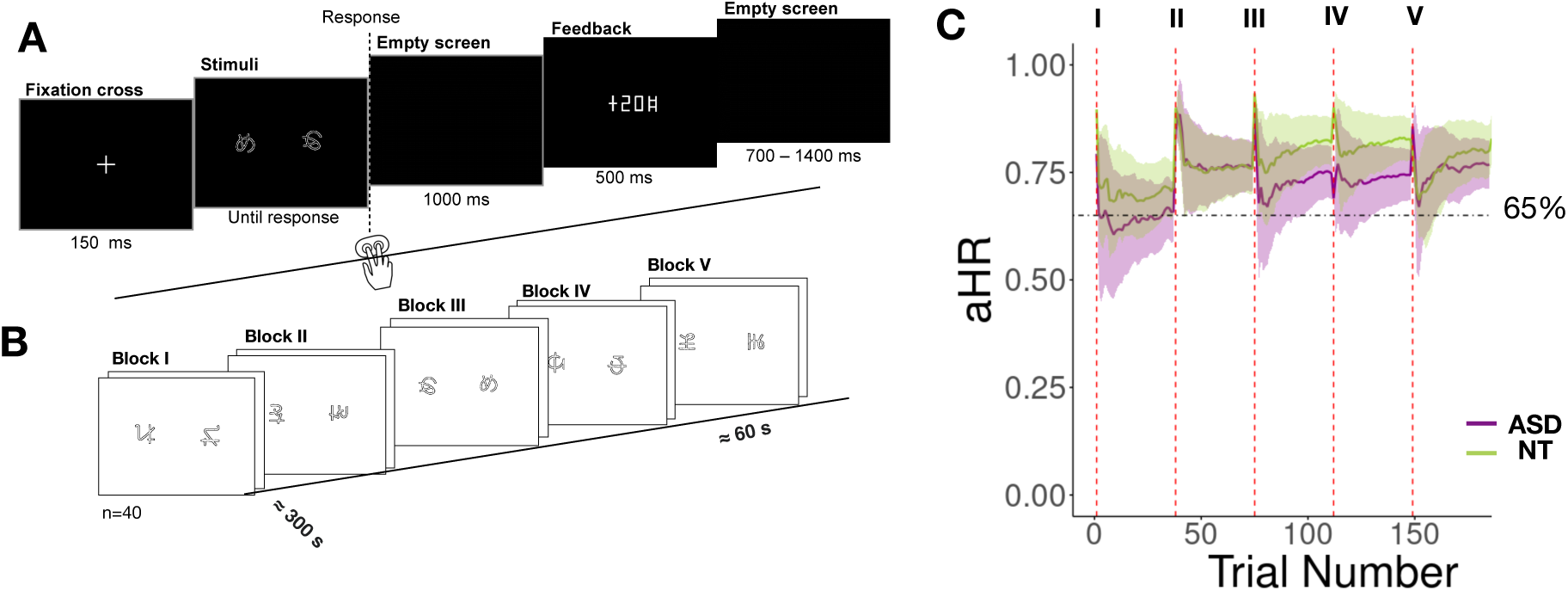
Probabilistic Two-Alternative Choice Task: Experimental Procedure and Behavioral Results. **(A)** Trial structure. **(B)** Timeline of the experimental procedure across blocks (black and white colors are inverted for clarity). The Roman numerals indicate the order of task blocks. **(C)** Cumulative probability of advantageous choices (advantageous choice hit rate, aHR) across trials within each sequential block for the NT (green) and ASD (purple) groups. Shaded areas represent the standard error of the mean (SEM). Vertical red lines indicate block onsets, and the horizontal dashed line marks the 65% aHR threshold.

Participants were presented with two stimuli simultaneously displayed on the screen and were asked to make a two-alternative choice in each block, with a new pair for each block. One stimulus in each block was associated with a higher probability of better outcomes (monetary gains or smaller losses compared to the other option on 70% of trials), resulting in a higher average payoff, referred to below as “advantageous stimuli” (AD). The other stimulus was associated with a higher probability of worse outcomes (monetary gains or smaller losses on 30% of trials), leading to lower average payoff, referred to below as “disadvantageous stimuli” (DA). The outcome probabilities remained constant within the blocks, and the sequence of gains and losses for the stimuli was generated quasi-randomly to prevent repeated outcomes in succession.

The probability of better and worse outcomes remained stable throughout the experiment, but the gain and loss magnitude changed between blocks to make them appear less similar and to encourage participants to learn during each block. The following magnitudes were applied: (1) −20/+20, (2) 0/+20, (3) −20/0, (4) +20/+50, and (5) −50/−20. The order of gain/loss magnitude was counterbalanced across participants, and three sequences were used: 1-2-3-4-5, 1-4-3-5-2, and 1-4-3-2-5. A new stimulus pair was used in each block.

Before the experiment, participants were informed that one stimulus was more advantageous than the other, but no additional information was provided. They were instructed to learn about the contingencies between their choices and outcomes through trial-and-error learning. Each trial began with a 150 ms fixation cross, followed by continuous presentation of the stimuli on the screen until the participant pressed a button (Fig. 1A). There was no time limit on the response time to reduce impulsive decisions. Participants’ responses were recorded using a handheld MEG-compatible fiber optic button response pad. Participants used the index and middle fingers of the right hand to select the stimulus appearing on the left and right side of the display, respectively.

Visual feedback was provided for 500 ms after each trial, indicating the number of points gained or lost on this trial. The points were accumulated throughout the experiment, and the current cumulative score was shown at the end of each block. The intertrial interval varied from 700 to 1,400 ms in a quasi-random order to keep the duration of the experiment relatively short and prevent fatigue and boredom (Fig. 1A).

The experiment comprised 40 trials in each of the five blocks (Fig. 1B). A short rest of about 1 minute (or longer if requested) was provided between blocks. The total duration of the experiment was approximately 35 minutes.

The Presentation 14.4 software (Neurobehavioral Systems, Inc., Albany, CA, United States) was used in the experiment.

### 2.3. Behavioral metrics

#### 2.3.1 Performance parameters

To assess how subjects generalize the rule associated with preference for AD stimuli, we calculated the advantageous choice hit rates (aHR), estimated as the ratio of AD choices to the total number of choices made. The aHR was calculated separately for each block (Fig. 1C) Also, the average hit rate of AD choices was estimated in each block.

Response time (RT) was measured from the stimulus onset to a button press. Trials with extreme RT values (< 300 ms and > 4,000 ms) were excluded from the analysis.

In addition, the “win-stay / lose-shift” metric was used to characterize the between-group differences in behavior. For trials with positive feedback (i.e., a ‘win’), we counted how often the participant chose the same stimulus in the next trial (a ‘stay’) versus choosing a different one (a ‘shift’). Similarly, we recorded how often participants avoided a stimulus following negative feedback (i.e., shifting after a loss). The number of occurrences for each behavioral strategy was summed and divided by the total number of trials in each block separately for each participant.

#### 2.3.2 Computational Model of Learning

We investigated how expectations (priors) were formed during learning in a stable probabilistic environment within a binary choice task. Priors were modeled as Gaussian probability distributions, characterized by their mean and precision (the inverse of variance). To model the decision-making process, we applied a two-level Hierarchical Gaussian Filter (HGF) using the pyHGF toolbox (Mathys et al., 2014).

The HGF framework captures how individuals update their beliefs based on observed outcomes. For each participant, the model received a sequence of 200 responses and their corresponding outcomes as input. The first level of the HGF represents trial-by-trial outcomes, while the second level reflects an individual’s beliefs about the probabilistic relationship between stimuli and outcomes, indicating the expected likelihood of encountering a particular outcome. To translate perceptual beliefs into decision-making, we paired the perceptual model with two alternative response models that map participants’ beliefs to their choices. Specifically, we used a response function based on (Shannon) surprise (Vossel et al., 2014). The response model quantifies how unexpected an outcome is, given the participant’s current belief state, with greater surprise leading to stronger belief updates. This mapping explains how beliefs guide choices – when an outcome aligns with expectations, belief adjustments are minimal, whereas unexpected outcomes drive greater adaptation.

The two-level HGF model was initialized with a temporary tonic volatility value of ω₂ = −4.0 at level 2, which was then optimized for each participant using a Bayesian framework implemented in PyMC3. The posterior distribution of ω₂ was sampled using Markov Chain Monte Carlo (MCMC), and the posterior mean was extracted as the final tonic volatility value for each participant. This optimized value replaced the initial temporary value, and the two-level HGF was re-run with the updated parameter.

We evaluated two key metrics: the second-level belief distribution (μ₂) and the second-level precision estimate (π₂). The belief μ₂ represents the probabilistic strength of a state being true, reflecting the expected initial value before incorporating observations. A higher μ₂ indicates a stronger initial belief about the hidden state. Expected uncertainty was quantified as the variance of the expectation, with its inverse termed precision (π₂). Higher prior precision corresponds to greater confidence in the prior belief.

### 2.6. Physiological data acquisition

MEG data were acquired using a 306-channel Elekta VectorView Neuromag array (Helsinki, Finland) with built-in filters (0.03–330 Hz) and a sampling frequency of 1000 Hz. The participants’ head shapes were measured by 3Space Isotrack II System (Fastrak Polhemus, Colchester, VA, United States) and digitized three anatomical landmark points (nasion, and left and right preauricular points) as well as approximately 60 randomly distributed points on the scalp.

We used two pairs of electrodes placed above and below the left eye and at the outer canthi of both eyes to record vertical and horizontal electrooculogram, correspondingly. We also recorded the bipolar electromyogram from the right neck muscles to identify muscular artifacts. For all recorded signals, the sampling rate was 1,000 Hz.

Pupil size recordings were obtained using the infrared eye tracker EyeLink 1000 Plus (SR Research Ltd., Canada). Details of the measurements and processing are provided in the companion paper by (Pultsina et al., 2024) Here, z-transformed pupil size data from the same participants were used to examine its relationship with measures of cortical activation.

### 2.7. MEG data preprocessing

We applied the temporal signal space separation method (tSSS) (Taulu et al., 2005) to the raw data using MaxFilter (Elekta Neuromag software). For further offline analysis, MNE-Python software was used (Gramfort et al., 2013). We removed cardiac artifacts and artifacts related to eye movements from continuous data using the fastICA method implemented in MNE-Python software. Specifically, we excluded contaminated epochs by thresholding the mean absolute signal values filtered above 70 Hz from each channel below 7 standard deviations of the across-channel average. Trials featuring extremely fast or slow response times (RTs) – specifically, those shorter than 300 ms or longer than 4,000 ms – were excluded from all analyses. MEG data were band-pass filtered at 1–40 Hz and resampled offline to 300 readings per second; this was safely above the Nyquist frequency since we were interested in MEG oscillations at lower frequencies not exceeding 30 Hz.

### 2.8. Trial selection

We primarily focused on the neural correlates of exploitative (model-congruent) and directed explorative (model-incongruent) choices, both of which can only be identified once a participant has acquired a predictive model (Pultsina et al., 2024). Therefore, individually for each participant and each block, we sampled exploitative and explorative choices using the following selection criteria. First, a participant made four consecutive advantageous choices without selecting a disadvantageous option. The probability of randomly achieving this sequence is 12.5% ((½)³ = 1/8). All trials preceding and including these four choices were excluded from the further analysis. Second, for the remaining trials in the block, at least 65% of choices had to be advantageous. This threshold was set to ensure that the performance significantly exceeded the chance level of 50%, as confirmed by a one-tailed one-sample binomial test (p < 0.05). Third, we defined an exploitative choice as an objectively advantageous choice surrounded by advantageous choices (eAD), and a directed explorative one as a disadvantageous one (eDA) surrounded by advantageous eAD choices. The third criterion was applied because our previous results revealed that the decision to explore a model-incongruent option affected physiological arousal in choices both preceding and following the directed explorative choice (Kozunova et al., 2022).

The MEG data were segmented into epochs from −1,750 to 2,750 ms relative to a response onset. The analysis included 2,821 eAD trials for the ASD group and 2,977 for the NT group. For eDA choices, 1227 trials from the ASD group and 1121 trials from the NT group were analyzed.

### 2.9. Source-level analysis and MEG signal processing

Participants underwent MRI scanning using a 1.5 T Philips Intera system (voxel size: 1 × 1 × 1 mm, T1-weighted images). Individual structural MRIs were used to reconstruct single-layer (inner skull) boundary-element models of cortical gray matter with a watershed segmentation algorithm (Ségonne et al., 2004) implemented in FreeSurfer 4.3 (Martinos Center for Biomedical Imaging, Charlestown, MA, USA). Anatomical surfaces were generated using the “recon-all” algorithm in FreeSurfer (Fischl et al., 2002) with default parameters. For 7 ASD and 2 NT participants, MRI data were not available; in these cases, the ‘fsaverage’ brain template from FreeSurfer was used. Head shapes were co-registered into a mesh using fiducial points and approximately 60 additional scalp points. A grid with a spacing of 4.9 mm was used for dipole placement, yielding 4098 vertices per hemisphere.

Time-frequency transformation and source localization were performed on single-trial epoch data. Data were transformed using the mne.minimum_norm.apply_inverse_tfr_epochs function in MNE-Python, which computes the time-frequency representation (TFR) of the epochs and applies the inverse operator to project these TFRs onto the cortical surface. Time-frequency analysis was performed using Morlet wavelets with a frequency resolution of 2 Hz and the number of cycles set to (frequency/2). Source reconstruction was achieved via sLoreta (Pascual-Marqui, 2002). The inverse solution was estimated using the following parameters: dipole orientation constrained to be orthogonal to the cortical surface (loose parameter set to 0.2), depth weighting parameter of 0.8, and signal-to-noise ratio set to 1. Preprocessed empty room recordings were used to estimate the noise covariance, which was applied during the inverse solution to account for sensor and environmental noise.

For our main analysis focusing on decision-making, we selected a time window from 900 to 300 ms before the decision termination (as indicated by a button press) to avoid potential overlap with effects related to visual stimulus onset and movement preparation (Tzagarakis et al., 2010). Within this time window, we concentrated on the α-β frequency band (10–30 Hz), based on previous studies showing power reduction in this range during working memory retrieval in decision-making (Hanslmayr et al., 2012). We therefore do not distinguish between α and β oscillations during the decision-making time window and refer to these oscillations as α-β from hereon. Visual inspection of time-frequency plots in cortical ROIs confirmed that this broad frequency range was most affected by decision-related processes in our task, supporting further analysis (Supplementary Fig. S2).

In an additional analysis, we evaluated the feedback-induced suppressive oscillatory response in the 200–600 ms post-feedback onset time window. This window was selected based on evidence of a strong reduction in EEG α-β power during this post-feedback time window (van Driel, 2012) and confirmed by our inspection of post-feedback time-frequency plots in the posterior cortical areas. For this analysis, only the α band (10-14 Hz) data were extracted and analyzed. This choice was made for two reasons: first, to assess the degree of visual attention to choice outcomes that is predominantly reflected by α suppression in the posterior cortex, and second, because beta-band responses to choice outcomes, particularly in anterior cortical regions, typically manifest as widespread synchronization rather than suppression – reflecting processes such as motor rebound or learning (Glazer et al., 2018; Pavlova et al., 2023; Pfurtscheller & Lopes Da Silva, 1999).

Baseline values were defined separately for each block using the stimulus-locked eAD epochs in the time window from −0.350 to −0.50 s relative to fixation cross onset. Power values for both baseline and response-locked epochs were log-transformed and multiplied by 10 for each participant and epoch independently. Subsequently, the averaged baseline values were subtracted from all time points, for each participant in each vertex and response-locked epochs, resulting in power values relative to the baseline expressed in decibels (dB) (Keil et al., 2022).

Baseline-normalized response power values in cortical sources were then averaged within cortical parcels defined by the HCP-MMP1.0 parcellation (Glasser et al., 2016) and morphed to the individual surface using ‘mne.morph_labels’ function. Time courses of these metrics were extracted using the ‘mne.extract_label_time_course’ function with the “mean” method and subsequently averaged across time points.

### 2.10 Statistical analysis

#### 2.10.1 ROI selection

As outlined in the Introduction, for the decision-making time window, we selected the following regions of interest (ROIs) based on previous literature on decision-making: the medial prefrontal cortex and anterior cingulate cortex (mPFC & ACC), the anterior insula and adjacent frontal operculum (AIC & FOP), and the ventral visual stream (VVS).

Initially, these ROIs were defined according to the HCP-MMP1 atlas (Glasser et al., 2016). Subsequently, the mPFC & ACC regions were refined based on the following considerations. The atlas delineates the anterior cingulate and medial prefrontal cortex into 15 areas along the medial aspect of each hemisphere, anterior to the motor and premotor regions. We excluded the five most ventral areas – the subgenual anterior cingulate cortex and four adjacent regions near the orbitofrontal cortex (OFC) – due to the limited sensitivity of MEG to OFC activity, likely influenced by the positioning of the sensor array (Ahlfors et al., 2010). The resulting mPFC & ACC boundaries closely align with those described by Dixon et al., 2017. Additionally, due to the limitations imposed by both mutual signal spread and signal cancellation between sources positioned along the interhemispheric fissure in the left and right hemispheres (Das et al., 2024; Hauk et al., 2022), we did not differentiate between the left and right mPFC and ACC in our analysis. The other ROIs – the anterior insula/frontal opercular region (AIC & FOP) and ventral visual stream (VVS) in each hemisphere – were defined within the boundaries specified in the HCP-MMP1 atlas.

For the feedback-related time window, ROI selection was guided by the observed spatial pattern of α power suppression, which spanned a broad swath of the posterior cortex in both hemispheres.

#### 2.10.2 Statistical analysis of MEG data

Statistical analysis was conducted using trial-wise linear mixed models (LMM) to examine oscillatory power suppression within the designated time windows. LMMs are well-suited for handling imbalanced designs, efficiently accommodate missing data without requiring listwise deletion, and are less sensitive to variations in trial numbers per condition compared to traditional ANOVA approaches based on grand averages (Kliegl et al., 2011).

We implemented LMMs using the **“lmer”** function from the **lme4** package in R (Bates et al., 2015). As an initial step, a parcel-wise one-sample *t*-test was performed to assess α-β power changes relative to baseline, with false discovery rate (FDR) correction applied across 360 parcels. Then, an LMM was applied to each of the 360 cortical parcels to examine the fixed effects of the factors of interest on α-β power suppression. However, in some parcels, the model failed to converge when the maximal random effect structure was included, resulting in the well-documented issue of model convergence (Park et al., 2020). To address this, we retained only the random intercept in the model, which may have inflated the probability of a Type I error (Barr et al., 2013).

To assess between-group differences in the effect of choice type (exploitative vs. explorative) during the decision-making time window, we applied the following LMM:

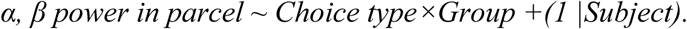

For the *p*-values associated with the interactions of interest, multiple comparisons correction was applied using the false discovery rate (FDR) method (Benjamini & Yekutieli, 2001) across 360 parcels. A significance threshold of 0.05 was applied to both unadjusted and FDR-corrected *p*-values.

For the final statistical analysis, power changes were averaged across parcels within the selected ROIs. In all subsequent analyses, the maximal random effect structure was specified, including random intercepts and slopes, to account for within-subject and within-item variability while ensuring model convergence.

For the ROI-level analysis of group differences in the effect of choice type on α- and β-power changes during the decision-making time window, we applied the following LMM:

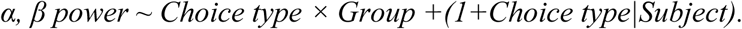

For effects following feedback onset, we used an LMM that included **Choice type, Feedback,** and **Group** as fixed effects, along with their interactions:

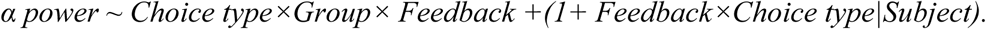

To assess the effect of block order on α- and β-power suppression associated with exploitative decisions (eAD) during the decision-making time window, as well as subsequent α suppression in the post-feedback time window, we implemented the following LMM:

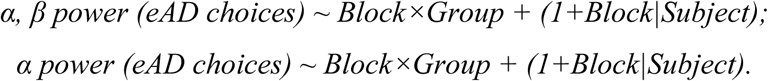

The estimated marginal means obtained from the LMM analyses were used both for generating visual representations and for conducting Tukey HSD post-hoc tests with the aid of the *emmeans* package in R (Searle et al., 2023).

#### 2.10.3 Statistical analysis of the behavioral results

To analyze the between-group difference in block-to-block dynamics of the RT and aHR, we used the following LMM:

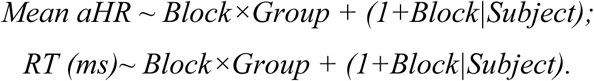

We also evaluated the effects of these factors and their interaction on the proportion between win-stay and lose-shift strategies using the following LMM model:

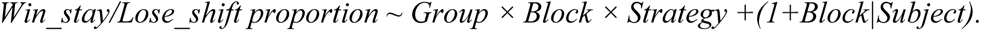

Trial-to-trial changes in precision and belief strength differences between groups were modeled using LMMs:

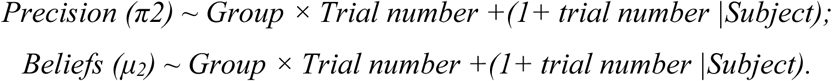

#### 2.10.4 Relation of α, β power changes to behavioral and autonomic measures

To determine whether the degree of α, β-power suppression during exploitative decisions influences later emerging responses to the decision made, such as reaction time (RT) and pupil dilation, we conducted single-trial regression analyses using the following LMM:

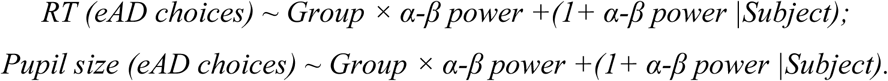

The z-transformed pupil size data for this analysis were extracted from the dataset described in the companion paper (Pultsina et al., 2024).

We also examined how trial-wise changes in precision and belief strength during exploitative (eAD) choices relate to decision-related α- and β-power suppression in the same trials:

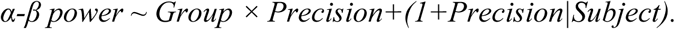

## 3 Results

### 3.1 Behavioral dynamics across blocks in ASD and NT adults

During our probabilistic learning task, participants learned to categorize choices between two stimuli in a pair as advantageous or disadvantageous ones based on a simple probabilistic classification rule. After the initial ‘learning’ block, they were expected to apply this rule to new, unfamiliar stimuli in subsequent blocks, which followed the same classification principle (Fig. 1C). Here, we analyzed whether rule generalization occurred, as reflected in performance changes across blocks, and whether there were differences in this process between the ASD and NT groups.

#### Performance parameters

For advantageous choice hit rates (aHR), we observed a positive order effect: a linear mixed model (LMM) revealed a significant effect of the Block factor (F(1,55.2) = 8.94, p = 0.004; Fig. 2A), indicating an increase in preference for advantageous choices from the first block to subsequent blocks (Tukey HSD: Block I vs. V, p = 0.004). A similar facilitating effect of block order was observed for response times (RT) in advantageous choices (F (1,19.1) =7.85, p = 0.011), with RTs decreasing significantly from the first block to later blocks (Tukey HSD: Block I vs. V, p = 0.011; Fig. 2B).

**Fig. 2.**
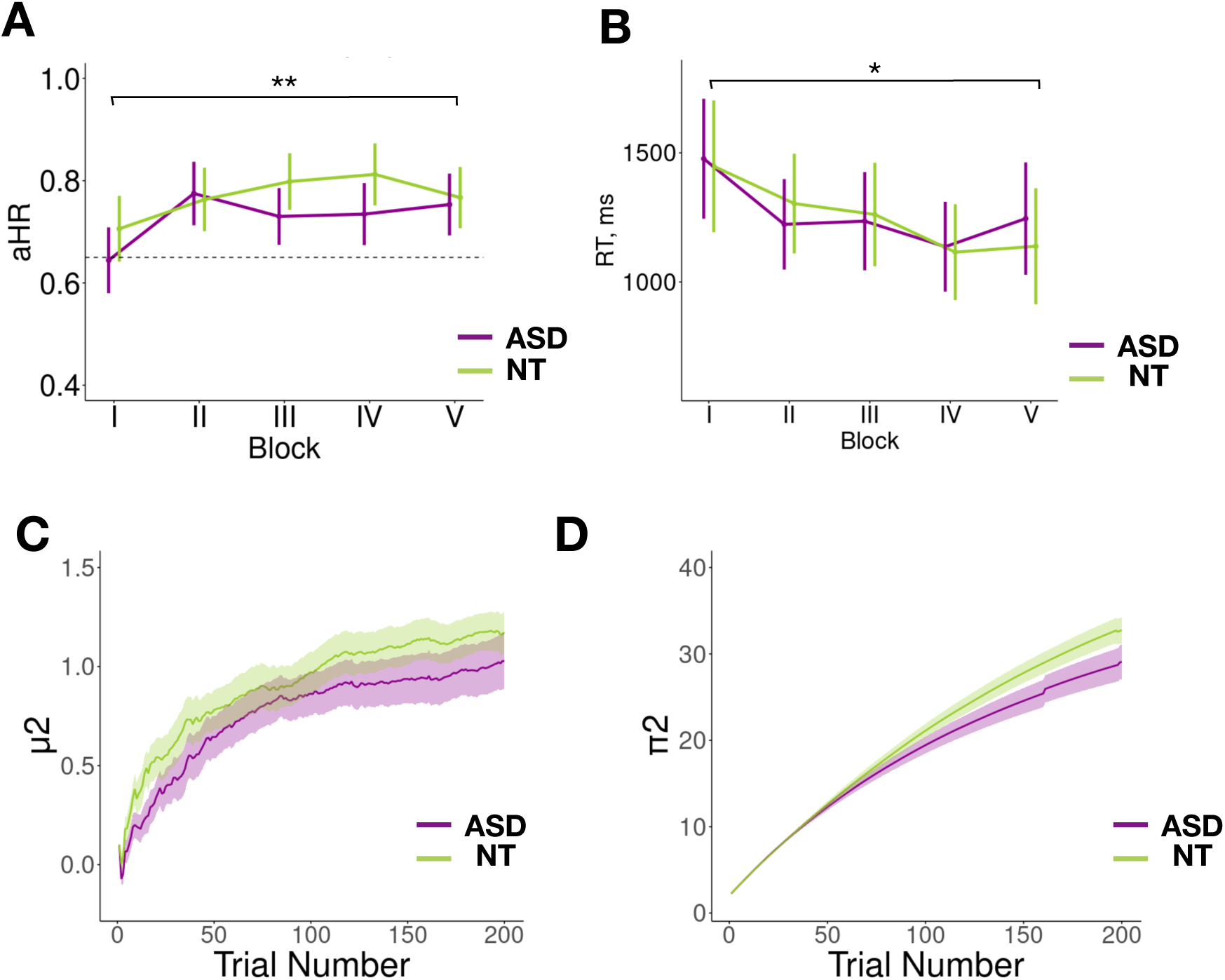
Behavioral dynamics in ASD and NT groups over the course of the task. **(A)** Block-order dynamics of the advantageous choice hit rate (aHR) and **(B)** response time for advantageous choices. The Roman numerals I–V indicate the position of each task block within the sequence. Points and error bars represent mean ± SEM. # p < 0.1, * p < 0.05 (LMM, Tukey HSD test). **(C)** Trial-wise dynamics of belief strength (mu2) and (D) belief precision (pi2). Shaded areas indicate the standard error of the mean (SEM) for each group.

However, we found no significant effects of group or interaction between group and block order for aHR: (Group: F(1,43.5) = 0.09, p = 0.768; Group*Block: F(1,43.3) = 0.006, p = 0.937). Similarly, there was no significant difference in the block-to-block dynamics of RTs (Group: F(1,28.3) = 0.56, p = 0.461; Group*Block: F(1,19.7) = 0.013, p = 0.912), as shown in Fig. 2B.

These findings suggest that, despite the novelty of stimulus pairs and win/loss values in each block, both ASD and NT participants recognized that the rule remained consistent across blocks and successfully applied it to maximize cumulative rewards. As a result, both groups showed improved accuracy and faster classification of new advantageous items in later blocks compared to the initial ‘learning’ block.

#### Beliefs strength and beliefs precision

We also modeled the “hidden parameters” of participants’ behavior using a two-level HGF. In the HGF, the individual trial-wise trajectories of the beliefs about the probabilistic mapping (HGF level i = 2) and informational or estimation uncertainty for level 2 are represented by their belief strength (μ^2^) and precision (π^2^).

The LMM revealed a gradual trial-by-trial increase in belief strength across both groups (Trial number: F (1,56) = 84.6, p < 0.001), with no significant between-group difference (Trial Number × Group: F (1,56) = 0.006, p = 0.939) (Fig. 2C).

Belief precision showed significant main effects of Trial Number (F(1,56) = 436, *p* < 0.001) and Group (F(1,56) = 7.36, *p* = 0.009), while the Trial Number × Group interaction (F(1,56) = 3.53, *p* = 0.066) did not reach statistical significance. Although belief precision increased across trials in both groups, this increase was attenuated in the ASD group (Fig. 2D). These findings suggest that, on average, individuals with ASD were less confident in their beliefs than neurotypical (NT) participants mainly because of slower confidence growth with task repetition.

While both groups showed a trial-by-trial increase in precision, this growth was smaller in the ASD group (Fig. 2D). These findings suggest that, on average, individuals with ASD were less confident in their beliefs than neurotypical (NT) participants, primarily due to slower confidence growth with task repetition.

In other words, participants with ASD were less precise in their prior beliefs compared to NT individuals, particularly in the second half of the experimental session (Fig. 2D), with minimal impact on performance. Lower belief precision may reflect greater variability in win-stay or lose-shift strategies, even if the overall proportion of these choices remained unchanged. To examine this possibility, we calculated the win-stay and lose-shift ratios, along with their variance, for each group (Fig. A1).

LMM revealed no significant effect of group (F(1,438) = 0.958, p = 0.328) or interaction between group and strategy (F(1,438) = 0.197, p = 0.657), consistent with comparable performance in ASD and NT individuals. In both groups, there was a significant effect of strategy (F(1,438) = 455, p<0.001) and its interaction with block (F(1,438) = 5.74, p=0.017). Both groups demonstrate a decrease in lose-shift behavior and an increase in win-stay behavior from block to block (Supplementary Fig. S1).

However, the win-stay ratio (but not the lose-shift ratio) exhibited significantly greater variance in the ASD group compared to NT participants (Fligner-Killeen test for between-group variance: χ² =8.05, p = 0.005), which may correspond to lower confidence in their beliefs.

### 3.2 Source level analysis of α-β suppression in the ROI

#### 3.2.1. Scaling of α-β suppression in exploitative versus explorative choices in ASD and NT individuals

Here, we tested the hypothesis that individuals with ASD over-engage brain resources when committing to an exploitative choice with a predictably advantageous outcome. As a result, their α-β suppression during decision-making – an index of resource allocation – may be comparable to confirmatory exploitative choices and typically more demanding explorative ones, which involve conflict and a high likelihood of unfavorable outcomes.

First, we determined the direction (suppression, enhancement, or baseline) of baseline-normalized α-β oscillation changes induced by decision-making across all subjects and choice types in the 360 cortical parcels defined by the HCP-MMP1 parcellation (see Methods). Supplementary Fig. S3 A, B illustrates widespread α-β suppression across the cortical surface, including the ROIs, where the reduction in oscillatory power primarily occurred within the α-β frequency range.

Next, we ensured that baseline α-β power did not differ significantly between ASD and NT participants in any cortical parcel (minimal p-value: df=44, t=1.941, p_fdr_=0.639; Supplementary Fig. S3). Thus, any group differences in decision-related power changes examined in the next step could not be attributed to baseline differences.

We initially applied LMM with a reduced random effect structure, which enhances sensitivity but also increases the risk of inflated false positives (see Methods for details), to explore the general spatial pattern of cortical parcels exhibiting significant Group × Choice Type interaction effects for α-β suppression. This analysis identified multiple cortical areas showing this effect, including the pre-selected ROIs – mPFC & ACC, AIC & FOP, and VVS (Fig. 3C).

**Fig. 3.**
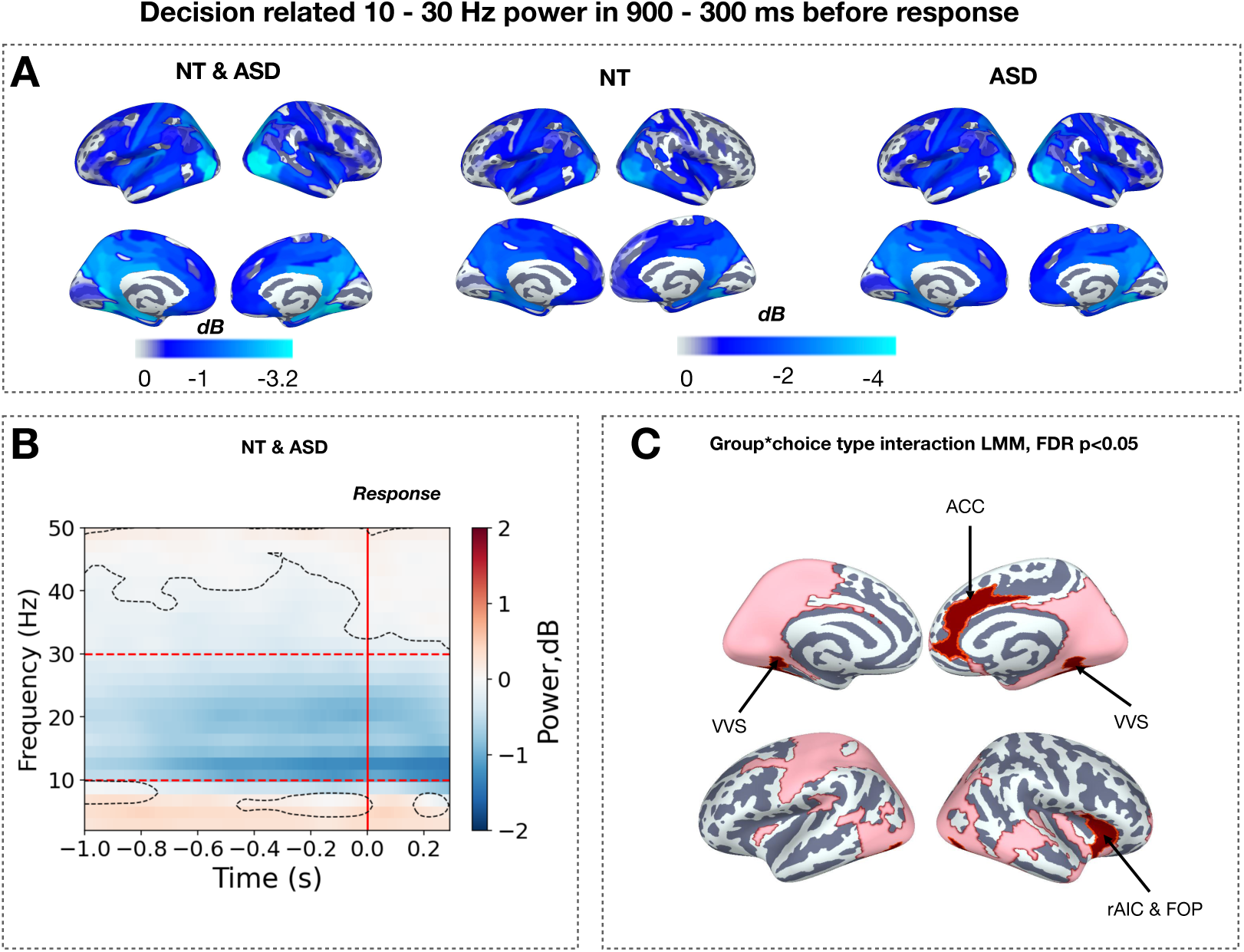
Decision-Related α-β Power Changes. **(A)** Cortical distribution of α-β power changes relative to baseline averaged across the decision-related time window (–900 ms to –300 ms before button press). Power changes are collapsed across all blocks and exploitative and explorative choice types and are shown for ASD and NT combined (ASD&NT), as well as for each group separately. Only significant cortical parcels are displayed (t-test, p < 0.001, FDR-corrected); **(B)** Time-frequency power changes relative to baseline, averaged across significant cortical parcels in ASD&NT sample. Curved dotted lines highlight frequency bands with significant power changes (t-test, p < 0.05, FDR-corrected). Red horizontal lines outline the selected α-β frequency range, and the vertical red line marks the termination of the decision period (button press); **(C)** Parcel-wise cortical distribution of Choice Type × Group interaction effects in α-β power change. Significant cortical parcels identified using LMM statistics (p < 0.05, FDR-corrected for 360 parcels) are shown in pink, while pre-selected regions of interest (ROI) are highlighted in maroon. Here and hereafter: mPFC&ACC - medial prefrontal and anterior cingulum areas, AIC&FOP - anterior insula, VVS – ventral visual stream areas.

To ensure robustness, we then performed an LMM analysis specifically within the ROIs, this time using the full random effect structure (Fig. 4).

**Fig. 4.**
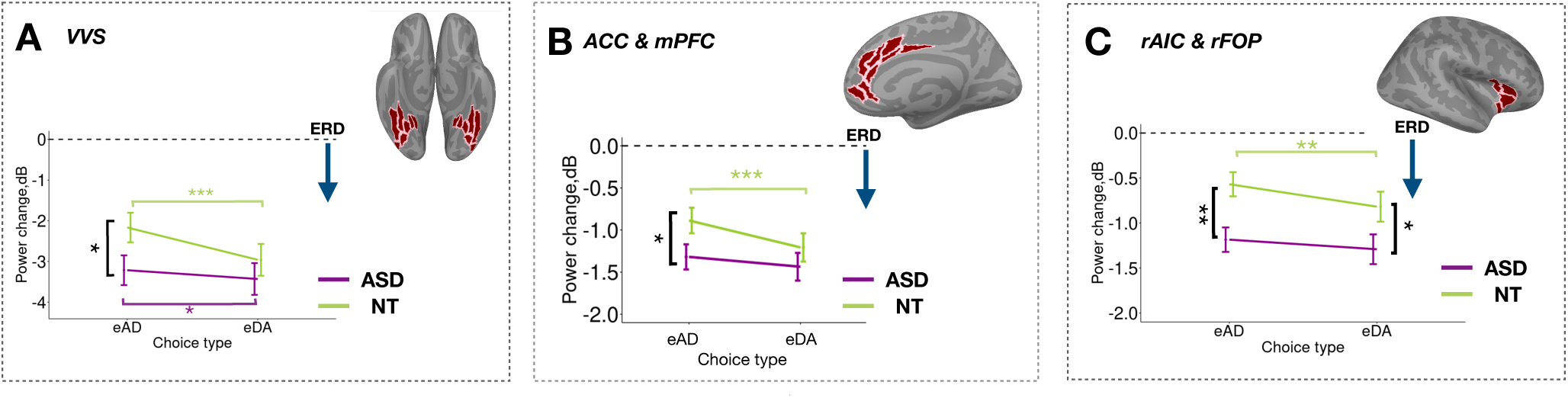
Decision-Related α-β Power Changes in ROIs During Explorative and Exploitative Choices in ASD and NT Adults. Points and error bars represent mean ± SEM of α-β power changes in the respective ROIs, averaged across all trials and subjects, as a function of choice type (exploitative [eAD] vs. explorative [eDA]) and group (ASD vs. NT). Tukey HSD test: *p < 0.05; **p < 0.005; ***p < 0.001. White lines delineate cortical parcels within the designated ROIs.

Significant Group × Choice type interactions were observed in the VVS (*F*(1,39.1) = 15.3, *p* < 0.001) and mPFC & ACC (*F*(1,437.9) = 7.44, *p* = 0.007). As shown in Fig. 4A and B, NT participants exhibited reduced α-β suppression for exploitative decisions compared to explorative ones (*Tukey HSD,* eAD vs eDA—VVS, *p* < 0.001; mPFC & ACC, *p* < 0.001). In contrast, this reduction was absent in ASD participants, who showed consistently high levels of α-β suppression regardless of decision type (*Tukey HSD,* eAD vs eDA—mPFC & ACC, *p* = 0.215; VVS, *p* = 0.045).

For the rAIC & FOP, there were significant main effects of Group (*F*(1,56.3) = 6.95, *p* = 0.011) and Choice type (*F*(1,42.8) = 9.9, *p* = 0.003), with no significant interaction (*F*(1,42.8) = 1.6, *p* = 0.214). This indicated that while α-β suppression in the right AIC was atypically strong in the ASD group for both decision types, it still showed a lower suppression for exploitative decisions compared to explorative ones, similar to the NT group (Fig. 4C).

No significant effect of factor group (*F*(1,39) = 0.530, *p* = 0.471) or its interaction with Choice type (*F*(1,32) = 1.96, *p* = 0.171) was present for the left AIC, and this ROI was excluded from all the further analyses.

These results suggest that, as expected, individuals with ASD lacked the typical ability to scale down α-β suppression when making decisions to commit to knowingly advantageous, preferred choices. An exception was the right anterior insula, where α-β suppression was relatively reduced for exploitative vs explorative choices but remained significantly higher in ASD compared to NT individuals (*Tukey HSD*, ASD vs NT: eAD, *p* = 0.002).

#### 3.2.2 Relationships between decision-related α, β power suppression with RT and pupil-linked autonomic arousal during exploitative trials

The relatively elevated α-β suppression during decision-making for preferred exploitative choices in ASD suggests increased deployment of brain resources for these easier decisions. To confirm this, we analyzed the relationship between trial-by-trial variations in α-β suppression within the cortical ROIs and changes in pupil-linked arousal and/or response time, which serve as behavioral measures of decision-making difficulty.

##### Pupil dilation

To examine the putative neuro-autonomic coupling, we extracted trials with exploitative choices and conducted trial-wise LMM regression with decision-related α-β suppression and Group as the predictors and pupil dilation as a response variable (Fig. 5 A,B). This analysis was done for each of the three ROIs separately.

**Fig. 5.**
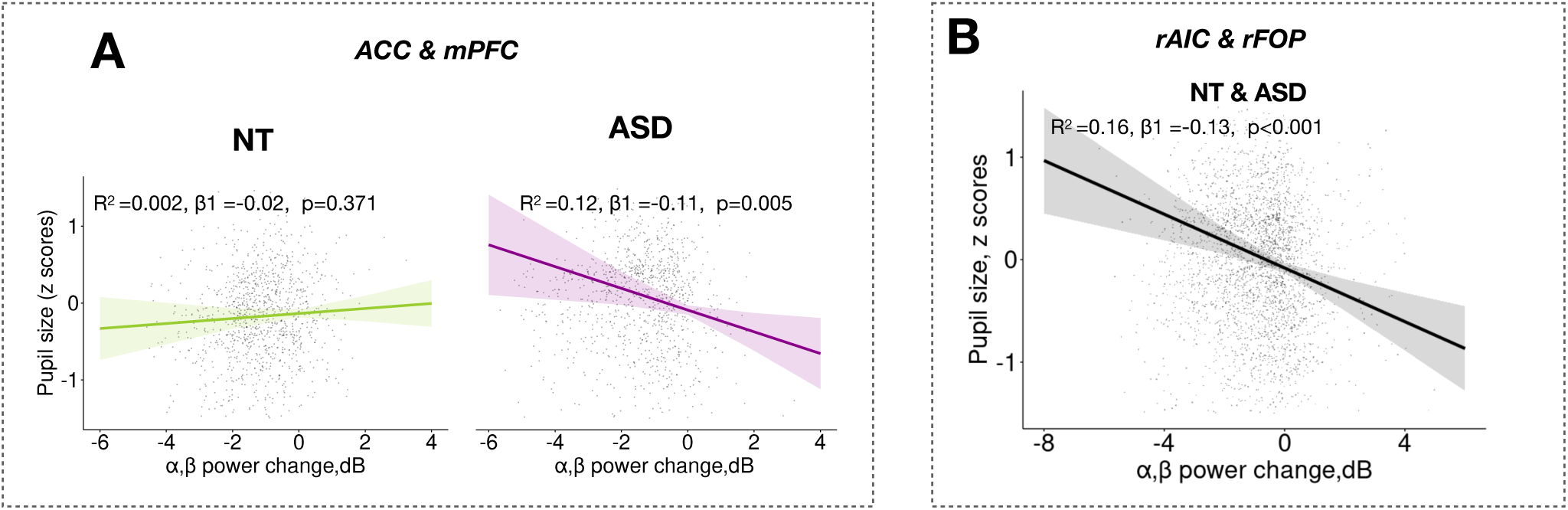
LMM regression of trial-wise pupil-linked arousal against decision-related α-β suppression during exploitative choices in ASD and NT participants. Only significant regression results are shown. The regression line for rAIC & rFOP is displayed for the pooled sample, as no significant difference in regression slopes was found between the NT and ASD groups.

The rAIC & rFOP was the main cortical region where functional activation, reflected in α-β suppression, reliably predicted the degree of pupil dilation in both groups, showing a directly proportional relationship with pupil-linked arousal (ASD & NT α, β power: *F*(1,41) = 14.1, *p*<0.001). In contrast, functional activation of the mPFC & ACC, which showed no significant relationship with pupil-linked arousal in neurotypical individuals, specifically predicted pupil dilation in the ASD group (Group × α, β power: *F*(1,38) = 4.34, *p* = 0.044; α, β power: NT: *F*(1,16.8) =0.847, *p* = 0.370, ASD: *F*(1,17) =10.2, *p* = 0.005). Lastly, activation in the VVS appeared to have no significant effect on pupil-linked arousal in either group (α, β power × Group: *F*(1,19) =0.581, *p* = 0.455; α, β power: NT: *F*(1,15) =0.02, *p* = 0.891, ASD: *F*(1,11) =0.652, *p* = 0.437).

Thus, in both groups, greater trial-wise decision-related activation of the right AIC &FOP predicted increased pupil-linked arousal following a preferred exploitative choice. Notably, in ASD, this association extended to the mPFC & ACC (Fig.5).

##### Response time

In addition to pupil-linked arousal, trial-by-trial functional activation in the cortical ROIs could also be related to the time spent on these relatively easy exploitative decisions. To examine this, we repeated the LMM regression analysis with decision-related α, β power change, and Group as predictors, and RT as the response variable for exploitative trials (Fig. 6). In both groups, we observed a strong, reliable relationship between α-β suppression and response time during decision-making, with greater suppression across all ROIs predicting longer response times: (mPFC & ACC: α-β power: F(1,31)=25.5, p<0.001, α-β power*Group effect F(1,31)=2.86, p=0.101; VVS: α, β power F(1,32)=101.5, p<0.001, α-β power *Group effect F(1,31.1)=0.67, p=0.419; rAIC & rFOP: α, β power F(1,27)=22.3, p<0.001, α-β power *Group effect F(1,27)=3.28, p=0.081).

**Fig. 6.**
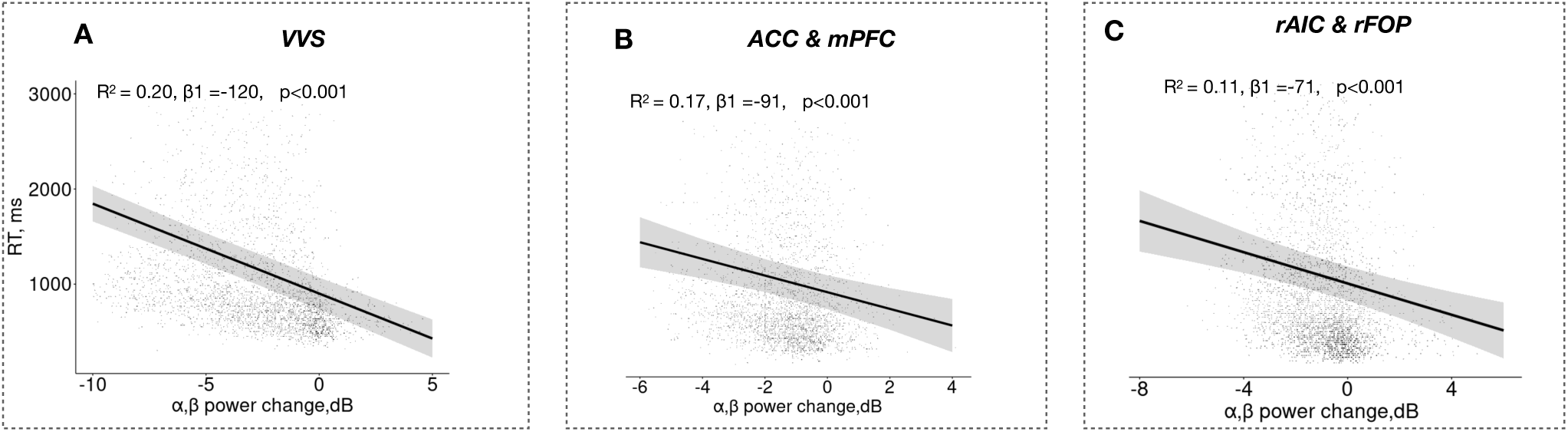
LMM regression of trial-wise response time against decision-related α-β suppression in ROIs during exploitative choices in the combined NT and ASD sample. The regression lines are displayed for the pooled sample, as no significant difference in regression slopes was found between the NT and ASD groups.

Taken together, these results suggest a robust link between **α-β suppression** and the allocation of processing resources during exploitative decision-making. In both NT and ASD groups, greater activation in the mPFC & ACC was most strongly associated with longer decision times, while right AIC & FOP activation was primarily linked to heightened phasic autonomic arousal during decision-making.

#### 3.2.3. Rule-based generalization: decision-related α-β power suppression changes across blocks

Previous findings suggest that heightened decision-related α-β suppression in ASD reflects an excessive allocation of brain resources to model-congruent exploitative choices. While individuals with ASD exhibited a typical increase in their preference for advantageous options after the initial learning block (Fig. 2A,C), the neural basis of this progression may differ from that of neurotypical individuals.

Fig. 7 shows that ASD vs. NT differences in decision-related α-β power gradually emerged after the first two blocks and persisted across the remaining three task blocks. LMM analysis comparing α-β power between the first and final blocks in the ASD and NT groups revealed a significant group × block order interaction for the mPFC & ACC region (F(1,35) =4.3, p = 0.046). While NT individuals exhibited a significant reduction in α-β suppression after the initial learning block (Tukey HSD, Block 1 vs. Block 5: NT, p = 0.023), this pattern was reversed in ASD participants, who instead showed increased activation toward the final block of the session (Tukey HSD, Block 1 vs. Block 5: ASD, p = 0.016). In the rAIC & rFOP, although the Group × Block interaction was not significant (F(1,30.3) = 0.50, p = 0.484), a significant main effect of Group indicated consistently higher rAIC & rFOP activation in ASD compared to NT participants across blocks (F(1,50.9) =10.1, p = 0.002; Fig. 7). In contrast, for decision-related α-β power changes in the VVS, neither the main effect of Group nor its interaction with Block order reached significance, though the trends pointed in the same direction (Group: F(1,50.3) = 3.45, p = 0.069; Group × Block: F(1,33.7) = 1.54, p = 0.223).

**Fig. 7.**
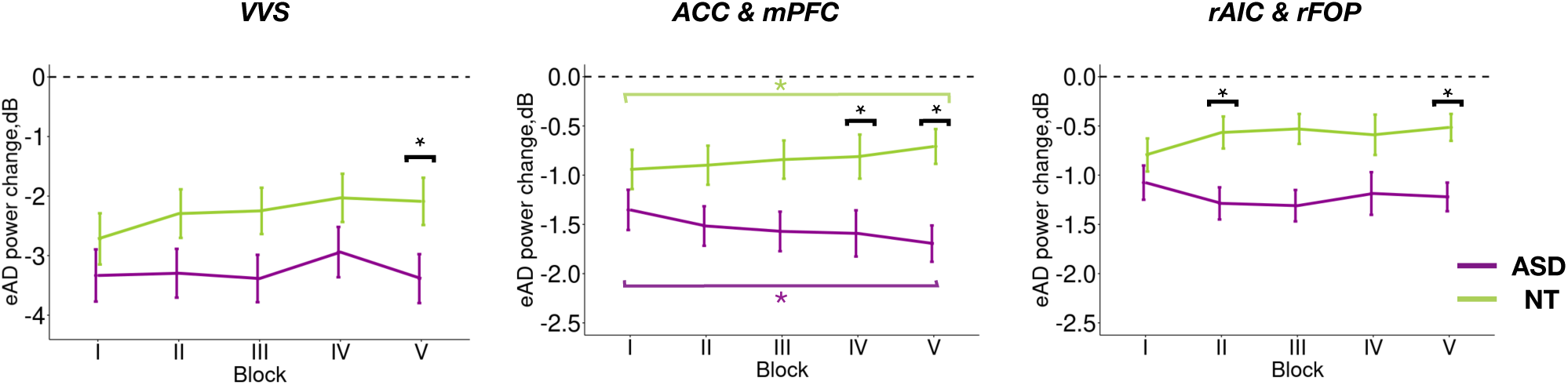
Difference in decision-related α-β power changes during exploitative choices between ASD and NT groups across blocks I to. **V.** All designations are as in Fig. 4.

These findings were unexpected, as behavioral measures in the ASD group – such as improved hit rates and faster response times – followed a typical trajectory of performance improvement across the block sequence, comparable to NT participants (see Fig. 2A,B). However, unlike NT individuals, this performance facilitation in ASD was accompanied by increased rather than reduced neural resource deployment.

This neural finding raised the question of whether α-β suppression is linked to the atypically low belief precision observed in ASD. Notably, belief precision was the only behavioral metric that differed between groups and followed a similar trajectory across blocks, with the gap between ASD and NT participants widening in the latter half of the session.

To directly examine this relationship, we conducted a trial-wise LMM regression analysis, using belief precision (π2) as a predictor and α-β suppression in the ROIs as the response variable (Fig. A4). In the NT group, higher precision was consistently associated with lower α-β suppression across cortical ROIs, indicating reduced cortical activation as precision increased (ACC & mPFC: F(1,17.1) = 4.58, p = 0.047; VVS: F(1,17.6) = 6.4, p = 0.021; rAIC & rFOP: F(1,19.3) = 6.98, p = 0.016). This suggests that greater confidence in beliefs typically facilitates the selection of probabilistically advantageous options, reducing cortical activation associated with decision-making.

In contrast, the ASD group showed no significant relationship between belief precision and α-β suppression in any ROI (ACC & mPFC: F(1,10.1) = 0.003, p = 0.956; VVS: F(1,12.8) = 0.44, p = 0.520; rAIC & rFOP: F(1,6.2) = 0.279, p = 0.615). Although the Group × Precision interaction effect was in the same direction across all three ROIs, it reached significance in the ACC & mPFC and rAIC & rFOP (ACC & mPFC: F(1,39.8) = 5.89, p = 0.020; rAIC & rFOP: F(1, 46.86) =6.6, p = 0.013), but not in VVS: F(1,44.2) =2.85, p = 0.098).

##### Feedback-related α-response

Although we mainly focused on the neural concomitants of a decision-making period, the feedback-related (FB) neural response was also of interest, as both HIPPEA and hypo-prior hypotheses predict increased attention to actual sensory evidence in ASD compared to NT participants. Additionally, if individuals with ASD overfit their internal models to observed data (as suggested by HIPPEA), one would expect a further enhancement in neural response to less likely outcomes – such as losses following advantageous exploitative choices or wins after typically disadvantageous explorative choices.

To test these predictions, we examined whether the feedback signal elicited stronger neural activation in individuals with ASD compared to NT participants and whether this activation was disproportionately greater for less likely versus more likely outcomes. For the FB-related neural response, we computed the mean α power change within the 200–600 ms time window following feedback onset for each cortical parcel, choice type, feedback type, and participant. As shown in Fig. 8A,B across all conditions, visual feedback primarily induced α suppression, which was most pronounced in posterior cortical regions, particularly the visual cortex (Fig. 8C). This finding is consistent with the well-established role of α suppression in enhancing attention to external visual signals.

**Fig. 8.**
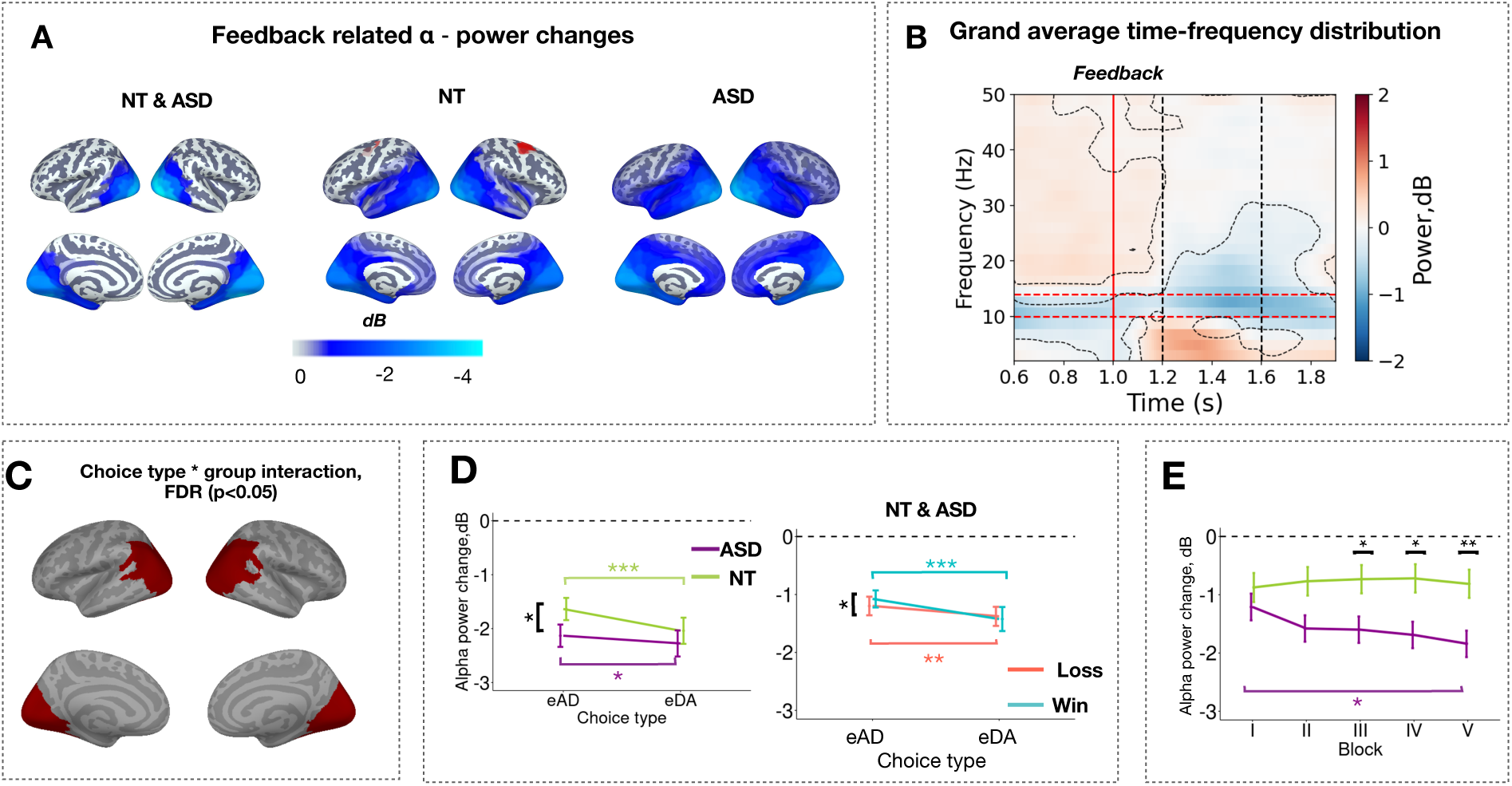
Feedback-related alpha power changes in NT and ASD groups. **(A)** Parcel-wise cortical distribution of α power changes relative to baseline for exploitative and exploratory choices combined, shown for all participants and separately for NT and ASD groups. Only significant parcels are displayed (one-sample t-test, p < 0.05, FDR-corrected); **(B)** Time-frequency analysis of power changes relative to baseline, averaged across all significant cortical parcels, choice types, and groups. The vertical red line marks feedback onset, and the dotted black lines indicate the feedback (FB) time window used for analysis. Curved dotted lines highlight frequency bands with significant power changes (t-test, p < 0.05, FDR-corrected). Red horizontal lines outline the α frequency band boundaries; **(C)** Parcel-wise cortical distribution selected ROI based on significant α power changes in the parieto-occipital cortex relative to baseline within the 200–600 ms post-feedback interval (p < 0.05, FDR-corrected for 360 parcels); **(D)** Feedback-related α power suppression, plotted separately by choice type (eAD, eDA: exploitative and explorative choices respectively) and group (left panel) and by choice type and feedback value (right panel).

We then performed an LMM analysis with Group, Choice Type, and Feedback (loss vs. win) as fixed factors, using a full random effect structure and mean feedback-related α suppression in the broad posterior cortical region as the dependent variable. This analysis revealed a significant effect of Choice Type (F(1,74.4) = 21, p < 0.001), illustrated in Fig. 8D. In both groups, feedback following rare explorative choices induced greater posterior α suppression than feedback after exploitative choices (Choice Type: F(1,143.1) =23.9, p < 0.001; Tukey HSD, eAD vs. eDA—NT: p<0.001, ASD: p = 0.042). The Group×Choice type interaction was also significant (F (1, 143.1) = 4.04, p = 0.046). As shown in Fig. 8D, the interaction effect was driven by stronger α suppression following feedback to exploitative choices in the ASD group compared to NT participants, suggesting their heightened attentiveness to actual choice outcomes regardless of choice type (Tukey HSD, eAD: NT vs. ASD, p = 0.027).

Additionally, a significant Choice Type × Feedback × Group interaction was observed (F(1,198.9) = 3.95, *p* = 0.048). As shown in Fig. 8D, this effect reflects that NT adults showed increased attention to negative feedback during exploitative choices—a pattern not observed in the ASD group (Tukey HSD, NT: eAD losses vs. gains, *p* = 0.002; ASD: *p* = 0.186). To further examine each group’s contribution to this interaction, we conducted separate linear mixed-effects model (LMM) analyses. These revealed a significant Choice Type × Feedback interaction in the NT group (F(1,74.5) = 4.5, *p* = 0.037), whereas no such interaction was found in the ASD group (F(1,106.7) = 0.03, *p* = 0.867).

The feedback results suggest that individuals with ASD exhibit a generally heightened attention-related neural response to probabilistic feedback following exploitative choices compared to NT participants, without a specific bias toward less likely outcomes.

## Discussion

Our findings in individuals with ASD reveal a dissociation between typical predictive behavior and atypical regulation of cortical resource allocation during probabilistic decision-making for the preferred, objectively optimal choices. This was observed in a value-based probabilistic task with a stable reward structure where one option consistently provided a probabilistically better outcome, despite changes in stimulus pairs across sequential blocks.

Behaviorally, both NT and ASD groups demonstrated increasing belief strength and precision – indicating growing confidence in their predictive model of the optimal choice – both within and across blocks, with belief strength approaching saturation by the middle of the block sequence (Fig. 2 C,D). This progression was accompanied by a growing bias toward the preferred choice and shorter decision times (Fig. 2B), suggesting that these exploitative decisions became easier as their high value was recognized with greater certainty through repetition. While ASD participants exhibited significantly lower confidence in their predictive model compared to NT participants – primarily due to a slower increase in the latter half of the block sequence – this did not significantly affect the dynamics of other behavioral parameters across blocks. These results indicate that high-functioning individuals with ASD demonstrated intact behavioral abilities to learn and generalize the probabilistic rule, according to which one alternative consistently provided higher rewards than the other, albeit with less confidence than NT participants.

Predictive abilities in high-functioning adults with ASD are sometimes described as suboptimal, particularly in volatile contexts like reversal learning tasks (Crawley et al., 2020; Sapey-Triomphe et al., 2022). In such environments, heightened state anxiety triggered by unpredictable contingency shifts may lead to both suboptimal behavior in ASD (Shi et al., 2022) and atypical neural processes underlying decision-making. However, in our probabilistic task with stable rules, we observed aberrant cortical activation patterns in ASD participants despite their preserved predictive behavior, eliminating predictive learning difficulties as the primary cause of these neural activity changes. Nevertheless, these findings do not exclude a potential link between atypically reduced confidence (belief precision) and altered cortical activation scaling.

In our study, decision-related cortical activation was assessed through the suppression of α-β oscillations in the anterior cingulate cortex, anterior insula, and ventral visual stream. α-β suppression was observed for both choice types: the preferred exploitative decisions, which aligned with experience-based predictions of favorable outcomes, and the less frequent deliberate directed explorative decisions, which conflicted with the predictive model and required greater efforts to commit (Pultsina et al., 2024). To investigate potential group differences in the load-dependent scaling of cortical resources, we relied on the prior evidence indicating that the degree of α-β suppression in task-relevant cortical regions reflects the neural resources allocated for information processing during tasks involving effortful memory retrieval or heightened attention to external stimuli (Barraza et al., 2023; Woodman et al., 2022).

We found that ASD participants displayed disproportionately high cortical resource expenditure during exploitative decisions – which may be perceived as easier, and which were associated with a lower likelihood of negative outcomes. This observation is supported by the following two ASD vs. NT comparisons.

The first comparison focused on exploitative versus explorative decisions. As reported in our previous study (Pultsina et al., 2024), both NT and ASD individuals showed shorter decision times and attenuated pupil-linked phasic arousal during exploitative decisions compared to explorative ones. In the current study, we observed that in NT participants, these model-congruent exploitative decisions also elicited weaker α-β power suppression across cortical areas of mPFC&ACC, right AIC & FOP, and VVS, likely indicating reduced cortical resource allocation for less effortful decisions (Fig. 4). The effort-dependent scaling of mPFC & ACC activity is consistent with findings from animal and human studies, which emphasize the critical role of the medial prefrontal cortex in effort-based decision-making, foraging behavior, and rule switching (Ma et al., 2019; for recent reviews see Clairis & Lopez-Persem, 2023; Klein-Flügge et al., 2022). Specifically, these regions are involved in an exploitation-exploration trade-off (Shenhav et al., 2016; Silvetti et al., 2018), which, in our task, is more prominent during switches to explore less frequently chosen options than during repeated exploitation of the preferred choice. The pronounced activation of the rAIC & rFOP during explorative rather than exploitative decisions (Fig. 4C) further support this interpretation, aligning with fMRI evidence linking AIC activation to strategy switching and increased task difficulty (Eckert et al., 2009; Wager & Barrett, 2004).

In contrast to the typical pattern, the ASD group exhibited comparably high activation across these cortical regions for both exploitative and explorative decisions (Fig. 4), suggesting a failure to adjust information processing resources appropriately. Rather than conserving neural resources for the less demanding exploitative choices, similar levels of information processing effort were engaged for both exploitative and explorative decisions.

Comparing this finding to previous literature is challenging, as the available brain studies on decision-making under uncertainty in ASD adults have primarily focused on post-feedback neural responses in the context of a reward prediction error (Cannon et al., 2021, for review; Carlisi et al., 2017; Mosner et al., 2019; Solomon et al., 2015). Notably, the only fMRI study examining the decision-making/anticipation phase of a probabilistic task (Iowa Gambling Task) in ASD versus NT participants reported findings similar to ours: a markedly reduced difference in prefrontal cortex activation between risky and safe choices, despite intact task performance (Carlisi et al., 2017). However, the comparative nature of previous findings does not clarify whether the abnormally subtle differential brain activation to risk as the source of increased uncertainty arises from cortical underactivation in both conditions (as the authors suggested) or from inflexible hyperactivation irrespective of the expected probability of choice outcome. Our MEG results rather suggest that in ASD, this effect reflects excessive neural resource allocation to decisions deemed more certain, although bearing some small degree of unavoidable uncertainty.

The idea that abnormally high cortical activation in ASD during exploitative decisions reflects increased mental effort is strongly supported by our trial-wise linear regression results (Fig. 5). Firstly, in both groups, greater α-β suppression in the mPFC & ACC and rAIC & rFOP during such decisions was associated with longer decision times. The link between increased prefrontal activation during relatively simple decisions and longer decision times and/or subjective decision costs aligns with prior neuroimaging findings (e.g., McGuire & Botvinick, 2010). Furthermore, in both ASD and NT participants, stronger α-β suppression in the right anterior insula reliably predicted phasic pupil-linked arousal elicited by exploitative choices (Fig. 5A). These non-invasive MEG findings complement rare evidence from human intracranial EEG studies demonstrating a causal link between task-evoked anterior insula activation and autonomic arousal, as measured by phasic pupillary dilation (Kucyi & Parvizi, 2020).

The second piece of evidence for excessive and inflexible neural resource deployment in ASD participants comes from the block-order dynamics of neural activation during exploitative decisions. As the task progressed, NT participants showed a significant reduction in α-β suppression during such decisions, particularly after the initial “learning” block (Fig. 6A). This suggests that rule generalization in NT individuals – applying a learned rule to optimize decisions with new stimulus pairs – not only improved decision speed and accuracy in later blocks (Fig. 2A,B), but also facilitated exploitative decisions. This facilitation likely reflected a shift toward a more “automatic” decision-making strategy, with reduced cognitive effort and emotional arousal, as indicated by decreased ACC and AIC activation. Such task practice effects, where improved performance requires fewer neural resources (Kelly & Garavan, 2005) are usually explained by increased neural efficiency.

In stark contrast to the typical dynamics of cortical activation associated with rule generalization, ASD participants exhibit comparable behavioral improvements, yet they sustain persistently high – or even further increased – decision-related activation in the mPFC & ACC and rAIC & rFOP across blocks (Fig. 6). This pattern suggests a failure to adjust neural resource allocation for exploitative decisions in proportion to the facilitative effects of rule generalization on behavioral performance. A similar pattern of persistently elevated cortical activation despite performance improvements was observed in an fMRI study on implicit visual pattern learning in ASD adults (Schipul & Just, 2016). Moreover, instead of the typical decrease in activation with practice, the study found increasing activation, suggesting a paradoxical escalation of effort allocation rather than a failure to downscale effort. We will revisit this point later.

Taken together, the comparisons examined in this study – exploration versus exploitation and initial versus later task blocks – indicate that individuals with ASD inflexibly allocate high brain resources during decisions involving predictably favorable yet uncertain outcomes. This occurs even though they clearly recognize the optimality of these decisions, as shown by their typical predictive behavior and ability to generalize rules with practice. Thus, while predictive ability appears largely intact in ASD, it may come at the cost of greater neural resource recruitment, even when cognitive demands are relatively low. Our neural findings provide stronger support for the “weak priors precision” hypothesis in ASD than the overly strong priors assumed by HIPPEA (Friston et al., 2013b; van de Cruys et al., 2014). If priors and their precision were excessively strong, brain resource allocation for model-congruent decisions would likely decrease, in contrast to the persistently high activation observed in our study.

Additionally, ASD participants exhibited heightened attention to sensory feedback, as reflected in greater feedback-related α suppression in the posterior cortex, regardless of choice type or feedback valence (Fig. 7D). This finding aligns with previous fMRI studies on reward prediction error (Mosner et al., 2019; Solomon et al., 2015). While both HIPPEA and the ‘weak priors precision’ hypothesis predict an exaggerated neural response to unsigned prediction error, HIPPEA’s proposed overfitting of internal predictions to observed evidence would also imply an abnormally heightened sensitivity to losses in model-congruent choices – an effect not observed in our ASD data (Fig. 7). Thus, our MEG findings reinforce the idea of relatively weak prior precision in ASD, consistent with previous autonomic findings in volatile environments (Lawson et al., 2017) and further demonstrate its impact on cortical activation patterns during a probabilistic task with a stable reward structure.

At the behavioral level, computational modeling directly identified atypically weak belief precision in our ASD participants, as indicated by abnormally low π², while belief strength (μ²) remained within the typical range (Fig. 2C). However, reduced belief precision alone does not explain the distinct trial-wise relationships between this computational parameter and decision-related cortical activation observed in NT and ASD participants (Supplementary Fig. S4). In NT individuals, higher precision was linked to lower cortical activation, reflecting a typical pattern where accumulated evidence reinforces confidence in model-congruent decisions, thereby reducing both the neural resources required and the time needed to reach a decision.

While this may appear inconsistent with recent EEG findings linking greater confidence in perceptual decisions to stronger posterior α suppression (Trajkovic et al., 2023), this discrepancy likely arises from task differences. In the Posner paradigm used by Trajkovic et al., 2023 decisions relied primarily on visual attention to a display, where lower attention—indexed by weaker posterior α suppression before and during stimulus presentation—was associated with reduced confidence in perceptual recognition. In contrast, our task primarily relied on memory retrieval, where increasing confidence in an optimal decision typically required fewer processing resources, leading to reduced α-β suppression in medial prefrontal and other cortical regions. These contrasting findings suggest that the relationship between cortical activation and subjective confidence may depend on the extent to which response selection relies on internal memory retrieval versus external perceptual processing.

Building on this reasoning, the absence of a significant relationship between confidence and cortical activation in ASD individuals during memory-based decision-making for exploitative choices, as revealed by trial-wise LMM regression, was unexpected. However, this finding is consistent with divergences in block-order dynamics across multiple measures. Unlike NT participants, who exhibit a coherent pattern, ASD individuals display a paradoxical trend: as their confidence in the objective advantages of exploitative choices increases across task blocks, their neural resource allocation to these decisions also increases rather than decreases (Fig. 7). Yet, similar to NT participants, this rise in confidence is still accompanied by faster decision times (Fig. 2B).

One plausible interpretation is that this disjointed pattern reflects a competing rather than cooperative interaction between rational and affective components of optimal decision-making in individuals with ASD. While task repetition and accumulated experience strengthened their confidence in the best possible outcome of exploitative choices, it also heightened their awareness that occasional losses were unavoidable due to inherent stochasticity. In ASD, this realization may carry heightened emotional salience, as indicated by increased pupil-linked arousal—a proxy for central arousal state—following exploitative choices, as reported in our companion paper (Pultsina et al., 2024). A gradual increase in negative affect alongside growing recognition of the prediction model’s utility could temper confidence growth as evidence accumulates, without weakening strong rational beliefs in the optimality of model-congruent choices. At the same time, it may counteract the typical link between increased confidence in prior beliefs and reduced cortical activation during model-congruent decisions, potentially explaining the paradoxical pattern observed in ASD. These findings suggest that belief precision, as inferred from behavioral performance, arises from the interplay of distinct neural processes that may either cooperate or compete during value-based decision-making, depending on task demands and clinical population.

Interestingly, the aversive response to irreducible errors may also explain findings from Schipul & Just, 2016, where cortical activation in ASD increased throughout a perceptual learning task, despite improved performance. That study set the initial recognition level at 70%, ensuring that errors occurred even as performance improved. It is possible that regardless of the specific experimental paradigm, the growing awareness of inevitable errors with task practice, combined with intense anticipatory emotion, may sustain or even amplify decision-related cortical activation in individuals with ASD. Furthermore, although this affective response to irreducible uncertainty did not disrupt predictive behavior in our simple probabilistic task, where the optimal choice followed a fixed rule, it may become a limiting factor in more complex learning environments, such as those with volatile conditions and shifting contingencies.

More broadly, while affective responses to high uncertainty are generally adaptive, aversion to the irreducible uncertainty of an action outcome in ASD may heighten the risk of avoidance behavior in any uncertain situation (Carleton et al., 2012). This well-documented “intolerance of uncertainty” in high-functioning adults with ASD (Hwang et al., 2020) was particularly pronounced in our sample (Pultsina et al., 2024), leading individuals to avoid even minimally uncertain situations, thereby limiting their opportunities for success in social environments (Solomon, 2020).

## Conclusion

In summary, high-functioning ASD individuals performing a two-alternative probabilistic task with unique stimulus pairs and a fixed optimal choice rule across five blocks exhibited typical rule generalization, improving decision speed and accuracy in later blocks. They also adjusted their prior beliefs and predictive model strength across trials and stimulus contexts, though with a notably smaller increase in precision values over time. Critically, unlike NT participants, rule generalization did not lead to a reduction in functional activation within decision-related cortical areas, suggesting a rigidly high allocation of neural resources despite increasing belief strength and confidence in model-congruent decisions. Additionally, on a trial-by-trial basis, increasing confidence was not associated with reduced cortical activation, as observed in NT individuals. We propose that these atypical neural-behavioral relationships in ASD may be due to heightened affective response to the awareness of inevitable losses, which are enhanced along with rational beliefs in the optimality of model-congruent choices. These findings suggest a maladaptive adjustment of neural resource allocation to expected uncertainty, rather than a fundamental impairment in probabilistic learning, underscoring the need for further empirical and computational studies to refine predictive coding models of ASD.

## Data Availability Statement

The code and raw data supporting the conclusions of this article are available by the authors upon reasonable request.

## Acknowledgments

We sincerely thank all the participants who participated in this study. The study was conducted at the unique research facility, Center for Neurocognitive Research (MEG-Center) of MSUPE.

## Author contributions

Pultsina Kristina: Conceptualization, Software & Formal analysis, Visualization, Writing - Original Draft & Review & Editing; Kozunova Galina: Methodology, Project administration, Writing - review & editing; Chernyshev Boris: Conceptualization, Writing - Review & Editing, Funding acquisition; Prokofiev Andrey: Project administration, Data Curation & recording; Tretyakova Vera: Software & Data Curation; Novikov Artem: Resources & Investigation; Rytikova Anna: Project administration & Data Curation; Stroganova Tatiana: Methodology, Conceptualization, Writing - Original Draft, Review & Editing, Formal analysis, Funding acquisition.

## Supplementary materials

**Supplementary Fig. S1.**
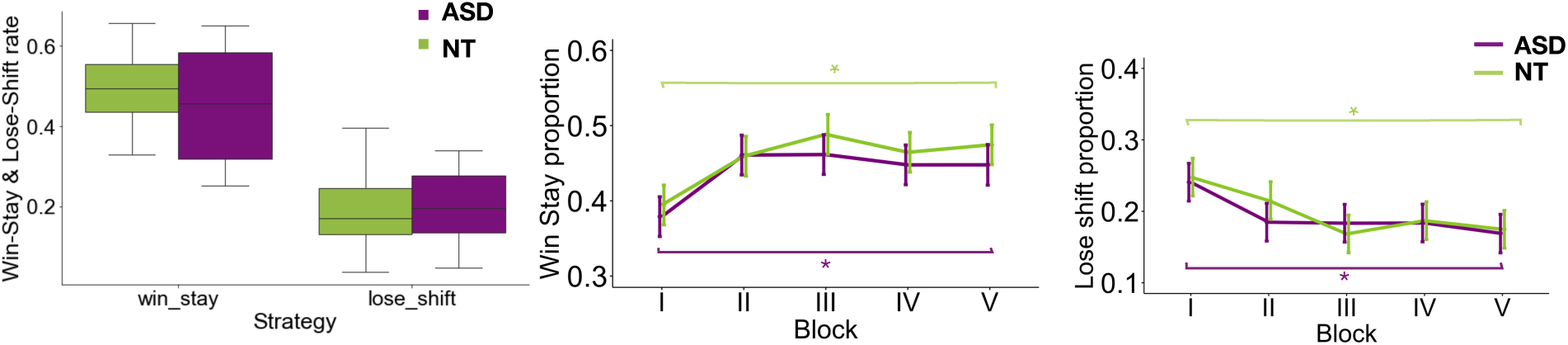
Win-stay and lose-shift proportion in ASD and NT groups.

**Supplementary Fig. S2.**
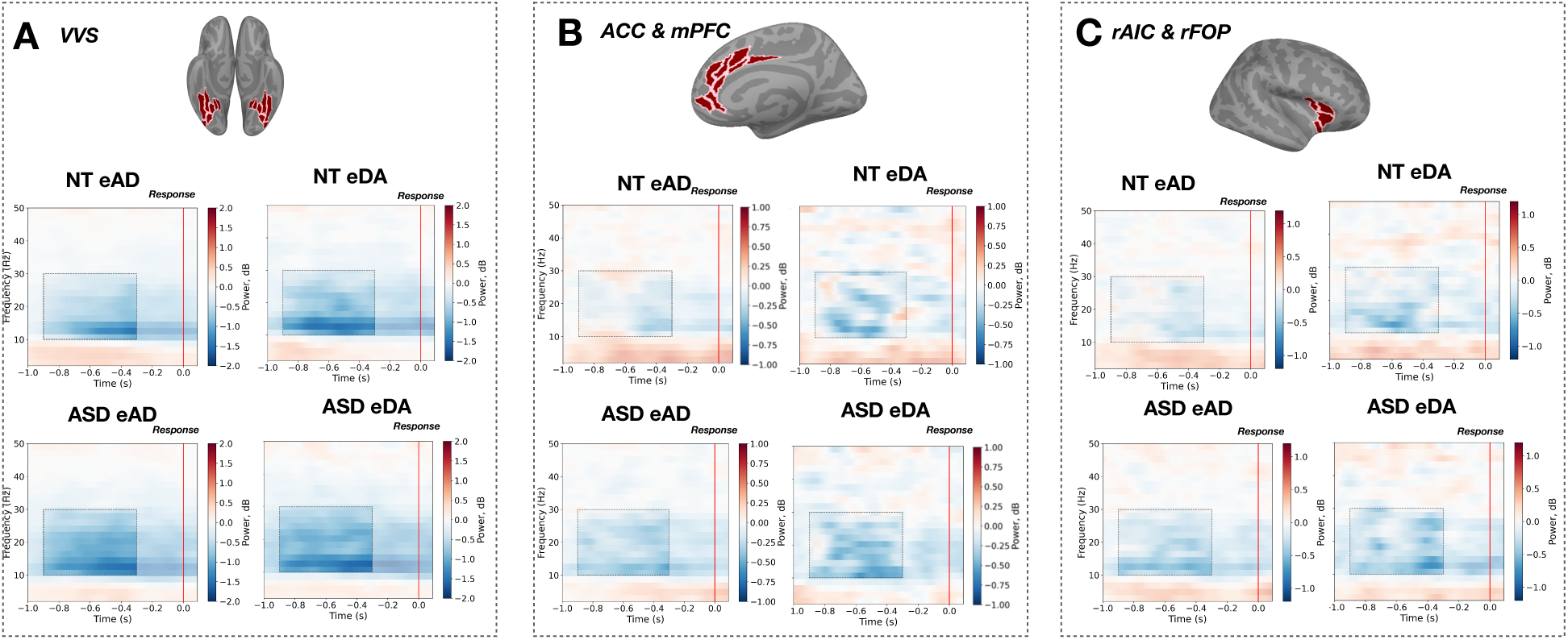
Time-frequency plots of cortical current changes in ROIs over the decision-making period during explorative and exploitative choice types in ASD and NT adults. eAD and eDA correspond to exploitative and explorative choices respectively. (**A) -** VVS – ventral visual stream. **(B)** – ACC & mPFC - anterior cingulate cortex and medial prefrontal cortex; (**C) –** rAIC & rFOP - right anterior insula.

**Supplementary Fig. S3.**
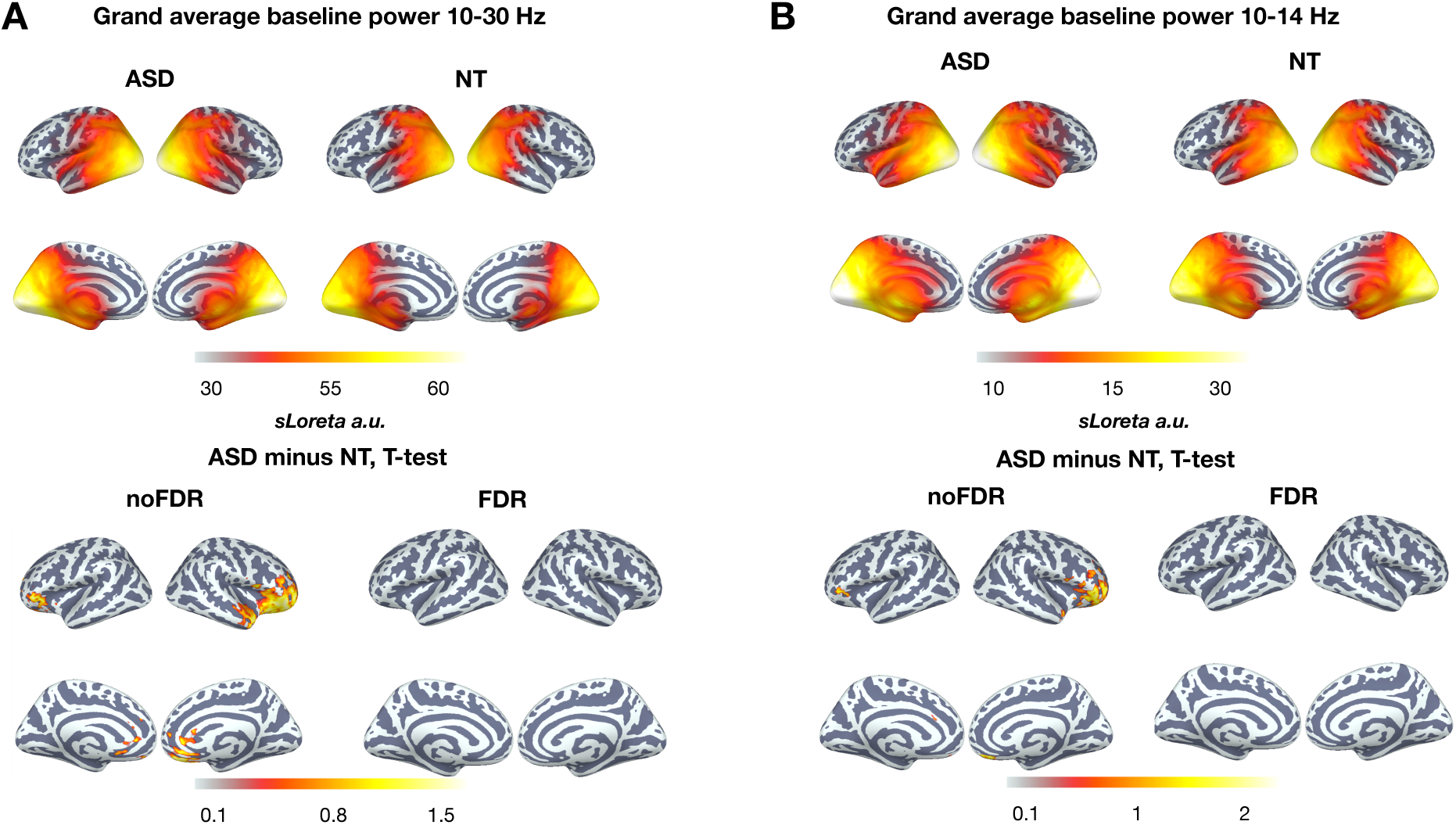
Comparison of Baseline Power Differences Between ASD and NT Groups. (A) Grand average baseline power in the 10–30 Hz frequency band (upper panel) and the corresponding differences between ASD and NT groups (lower panel). Only significant cortical parcels are displayed, both with and without FDR correction (p < 0.05). (B) Grand average baseline power in the 10–14 Hz frequency band and its differences between ASD and NT groups. Only significant vertices are shown, both with and without FDR correction (p < 0.05).

**Supplementary Fig. S4.**
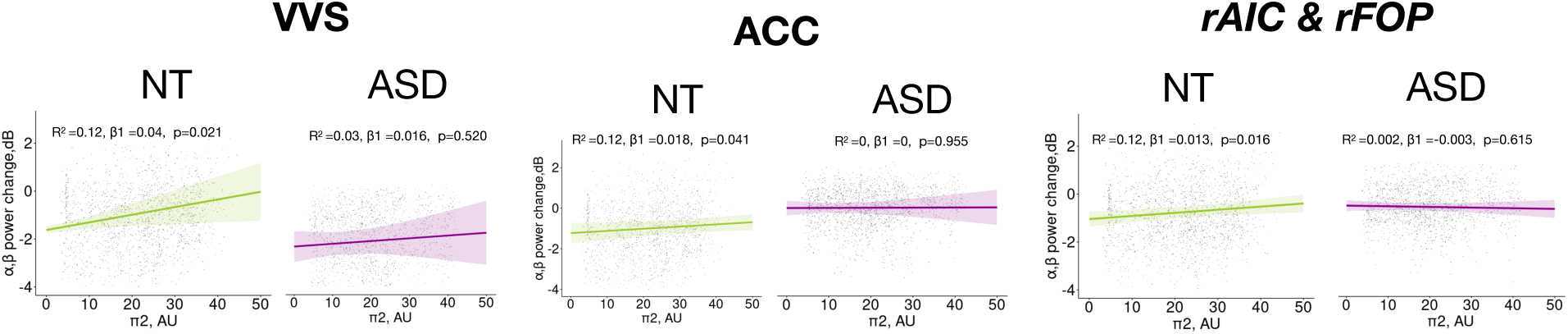
Trial-wise LMM regression of decision-related α-β suppression during exploitative choices as a function of prior precision in ASD and NT groups.

