## Appendix for "Neural Adaptation to Expected Uncertainty in Neurotypical Adults and High-Functioning Adults with Autism Spectrum Disorder"

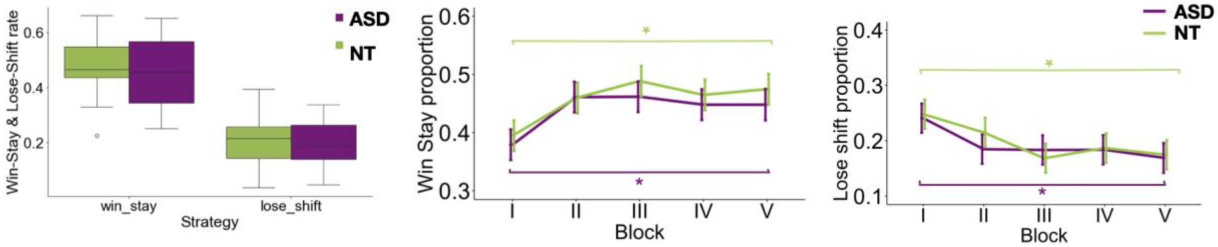


**Figure A1.** Win-stay and lose-shift proportion in ASD and NT groups.

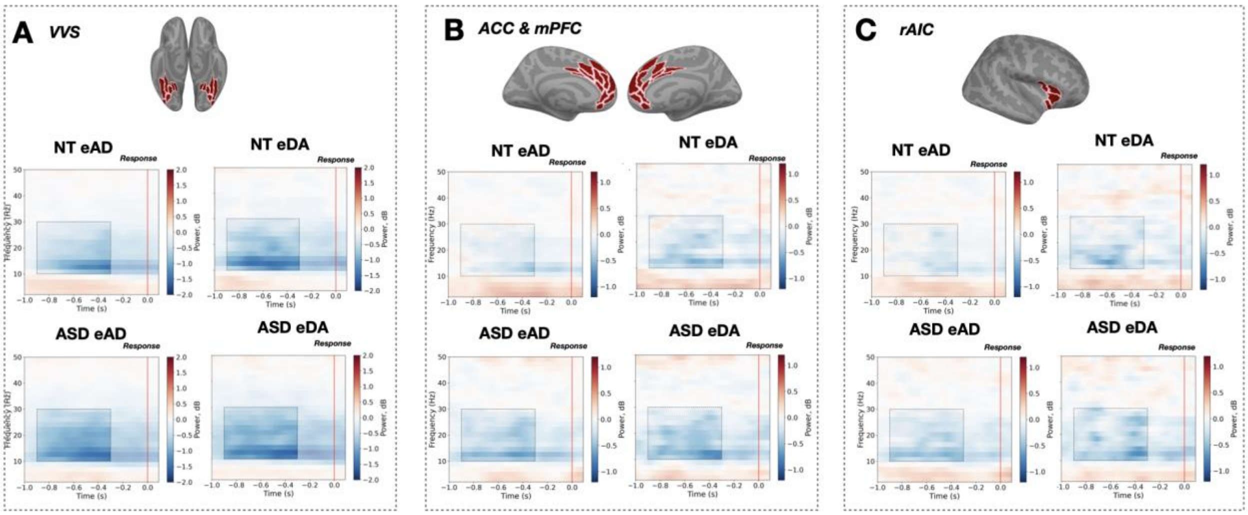


**Figure A2.** Time-frequency plots of cortical current changes in ROIs over the decision-making period during explorative and exploitative choice types in ASD and NT adults. eAD and eDA correspond to exploitative and explorative choices respectively. (**A) -** *VVS* – ventral visual stream. **(B)** – *ACC & mPFC* - anterior cingulate cortex and medial prefrontal cortex; (**C)** – *rAIC & rFOP* - right anterior insula.

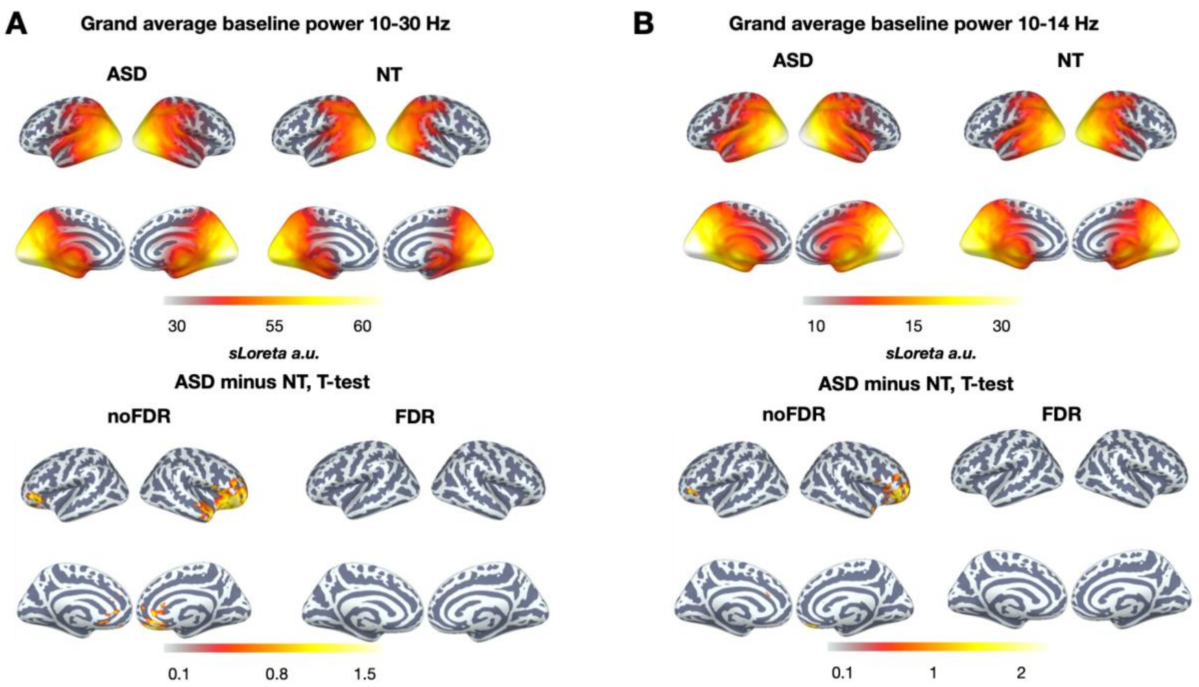


**Figure A3.** Comparison of Baseline Power Differences Between ASD and NT Groups.

(A) Grand average baseline power in the 10–30 Hz frequency band (upper panel) and the corresponding differences between ASD and NT groups (lower panel). Only significant cortical parcels are displayed, both with and without FDR correction ($p < 0.05$).

(B) Grand average baseline power in the 10–14 Hz frequency band and its differences between ASD and NT groups. Only significant vertices are shown, both with and without FDR correction ($p < 0.05$).

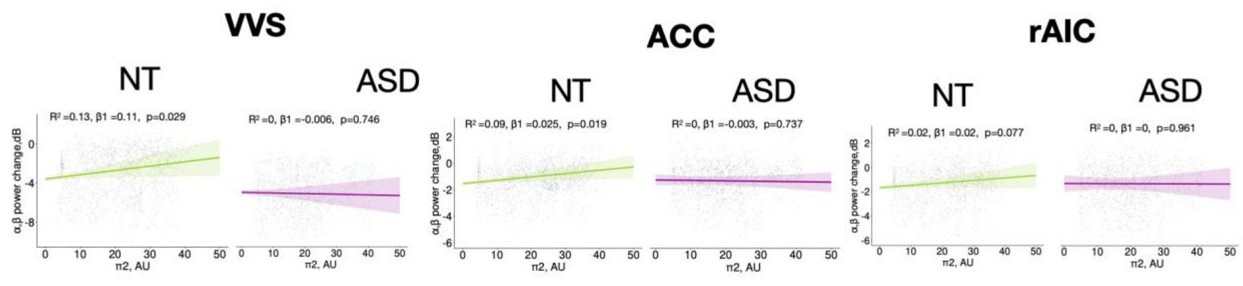


**Figure A4.** Trial-wise LMM regression of decision-related α-β suppression during exploitative choices as a function of prior precision in ASD and NT groups.
